# Towards robust and replicable sex differences in the intrinsic brain function of autism

**DOI:** 10.1101/2020.06.09.142471

**Authors:** Dorothea L. Floris, José O. A. Filho, Meng-Chuan Lai, Steve Giavasis, Marianne Oldehinkel, Maarten Mennes, Tony Charman, Julian Tillmann, Guillaume Dumas, Christine Ecker, Flavio Dell’Acqua, Tobias Banaschewski, Carolin Moessnang, Simon Baron-Cohen, Sarah Durston, Eva Loth, Declan G. M. Murphy, Jan K. Buitelaar, Christian F. Beckmann, Michael P. Milham, Adriana Di Martino

**Author notes:** **Corresponding author:** Adriana Di Martino, MD Child Mind Institute, Autism Center, 101 E 56 Street, NY, NY 10026. **equal contribution** (Dorothea L. Floris, José O. A. Filho).

## Abstract

**Background:** Marked sex differences in autism prevalence accentuate the need to understand the role of biological sex-related factors in autism. Efforts to unravel sex differences in the brain organization of autism have, however, been challenged by the limited availability of female data.

**Methods:** We addressed this gap by using a large sample of males and females with autism and neurotypical (NT) control individuals (ABIDE; Autism: 362 males, 82 females; NT: 409 males, 166 females; 7-18 years). Discovery analyses examined main effects of diagnosis, sex and their interaction across five resting-state fMRI (R-fMRI) metrics (voxel-level Z > 3.1, cluster-level P < 0.01, gaussian random field corrected). Secondary analyses assessed the robustness of the results to different pre-processing approaches and their replicability in two independent samples: the EU-AIMS Longitudinal European Autism Project (LEAP) and the Gender Explorations of Neurogenetics and Development to Advance Autism Research (GENDAAR).

**Results:** Discovery analyses in ABIDE revealed significant main effects across the intrinsic functional connectivity (iFC) of the posterior cingulate cortex, regional homogeneity and voxel-mirrored homotopic connectivity (VMHC) in several cortical regions, largely converging in the default network midline. Sex-by-diagnosis interactions were confined to the dorsolateral occipital cortex, with reduced VMHC in females with autism. All findings were robust to different pre-processing steps. Replicability in independent samples varied by R-fMRI measures and effects with the targeted sex-by-diagnosis interaction being replicated in the larger of the two replication samples – EU-AIMS LEAP.

**Limitations:** Given the lack of *a priori* harmonization among the discovery and replication datasets available to date, sample-related variation remained and may have affected replicability.

**Conclusions:** Atypical cross-hemispheric interactions are neurobiologically relevant to autism. They likely result from the combination of sex-dependent and sex-independent factors with a differential effect across functional cortical networks. Systematic assessments of the factors contributing to replicability are needed and necessitate coordinated large-scale data collection across studies.

## Background

Autism spectrum disorder (autism) is characterized by a marked male preponderance in prevalence with three times more males being diagnosed than females [1]. This pronounced sex-differential prevalence implies that sex-related biological factors are likely implicated in the neurobiology of autism. However, little is known about the differential underlying neural expressions in males and females with autism. Such knowledge could widen our understanding of potential underlying mechanisms of autism and related neurodevelopmental conditions [2].

This has motivated research into the impact of biological sex on brain organization in autism [2–5]. With the widely accepted view that the neurobiology of autism involves differences in large-scale brain networks [6,7], resting-state functional magnetic resonance imaging (R-fMRI) has proven to be a valuable complementary tool for investigating atypicalities in intrinsic functional connectivity (iFC). While the exact nature of the intrinsic brain organization in autism remains to be established [6], research on the impact of biological sex differences in autism is just beginning to emerge.

Several R-fMRI studies have focused on autism-related sex differences in iFC [2,8,17,18,9–16]. They vary on the extent of the functional networks and intrinsic properties examined. Most of them examined the strength of iFC between one or more regions/networks selected *a priori* [8–10,12,13,17,18], or via data-driven analyses [16]. A few others investigated either local or homotopic iFC across the whole brain [2,11,15]. Across these different efforts, the pattern of findings have also been mixed; some studies supported the predictions from the ‘extreme male brain theory’ [10,11], others supported the predictions from the ‘gender-incoherence’ theory [8,9,12,13,16]. The extreme male brain theory model predicts that brain characteristics in males and females with autism will resemble those in neurotypical males (i.e., shifts towards maleness in both sexes [19]). R-fMRI results consistent with a shift towards maleness in autism were reported in both Ypma et al. [10] and Kozhemiako et al. [11,15]. The ‘gender incoherence’ model predicts that brain characteristics in females with autism resemble those of neurotypical males, whereas brain characteristics in males with autism resemble those of neurotypical females (i.e., androgynous patterns in the sexes [20]). The ‘gender incoherence’ model has been supported by findings from prior R-fMRI studies [8,9,12], where the results largely revealed hyper-connectivity in females with autism similar to neurotypical (NT) males and hypo-connectivity in males with autism similar to NT females. Such seemingly inconsistent findings of sex-related differences were in part addressed by Floris et al. [2] who showed that, at least in males with autism, distinct patterns of atypical sex-differentiation coexist, and vary as a function of the neural networks involved. However, the intrinsic brain organization in females with autism has remained largely unclear and the scarce availability of female datasets in most studies may have contributed to the variability in findings in males and females [21,22].

Accordingly, to explore sex-related atypicalities in autism relative to NT controls, we used, as discovery sample, a large R-fMRI datasets of both males and females of autism and NT selected from the Autism Brain Imaging Data Sharing Exchange (ABIDE) [22,23]. By aggregating neuroimaging datasets from multiple sources, this data sharing initiative has begun to provide a means to address the challenge of underrepresentation of female datasets in autism research. Examining both sexes in both autism and controls allows to directly capture not only sex differences that are common across individuals (i.e., regardless of their diagnosis [main effect of sex]), but also those that are specific to autism and point towards atypical autism-specific sex differential patterns (i.e., sex-by-diagnosis interaction effects) [4]. To do so, given prior inconsistencies in the literature and the limited insights onto the brain organization of females with autism, we used a discovery approach. Unlike most prior work that focused on specific networks or circuits selected *a priori,* we investigated the whole-brain across multiple R-fMRI metrics. We selected R-fMRI metrics capturing unique aspects of the intrinsic brain organization during typical development [24,25] and, most germane to this study, being reported to be involved in typical sex differences and be affected by autism. They comprised: 1) posterior cingulate cortex (PCC)-iFC – e.g., [2,10,23,26–32]; 2) voxel-mirrored homotopic connectivity (VMHC) [33] – e.g., [11,32–35]; 3) regional homogeneity (ReHo) [36] – e.g., [15,32,37,38]; 4) network degree centrality (DC) [39] – e.g., [32,39–41]; and 5) fractional amplitude of low frequency fluctuations (fALFF) [23,42] – e.g., [23,32,43].

Beside the role of small female samples, prior inconsistencies in autism-related sex differences in R-fMRI can be due to other factors that can impact reproducibility. For example, while there are growing concerns on the role of pre-processing strategies [44,45], a recent study showed that their impact on autism-related mean group-differences is minimal [46]. Additionally, while several studies have reported some degree of consistency on R-fMRI findings across either independent, or partially overlapping, samples [41,47–49], results from other studies have raised concerns on the replicability of group-mean diagnostic effects [46,50]. However, none of these studies have explicitly examined robustness and replicability of sex-by-diagnosis interaction effects, which account for a potentially relevant source of variability in autism – biological sex. Thus, we conducted secondary analyses to assess the extent to which the pattern of findings obtained in our discovery analyses were also observed *a)* after applying different nuisance pre-processing steps that have been previously validated, though used inconsistently in the autism literature [46], and *b)* across two independent, multisite R-fMRI datasets: the EU-AIMS Longitudinal European Autism Project (LEAP) [51,52] and the Gender Explorations of Neurogenetics and Development to Advance Autism Research (GENDAAR) dataset [53] – i.e., *robustness* and *replicability*.

## Methods

### Discovery sample: ABIDE I and II

For discovery analyses, we examined the R-fMRI dataset with one of the largest number of females and males in both the autism and the NT groups available to date, selected from the Autism Brain Imaging Data Exchange (ABIDE) repositories ABIDE I and II [22,23]. The final ABIDE I and II dataset of N=1,019 included N=82 females with autism, N=362 males with autism, N=166 neurotypical females (NT F), and N=409 neurotypical males (NT M), aggregated across 13 sites. Specific selection criteria are described in Supplementary Material in the Additional file 1 and depicted as a figure in the Additional file 2. Briefly, we selected cases between 7-18 years of age (the ages most represented across ABIDE sites), with MRI data successfully completing brain image co-registration and transformation to standard space, with FIQ between 70-148 and with mean framewise displacement (mFD) [54] within three times the interquartile range (IQR) + the third quartile (Q3) of the sample (i.e., 0.39 mFD). Further steps included matching for mean age *across* groups as well as for mFD and IQ *within* diagnostic groups. This latter step limited the number of exclusions while keeping average group motion low (mFD<.2mm) and sampling biases that may result when matching neurodevelopmental conditions to NT around intrinsic features such as IQ [55,56]. At each step, any sites with less than three individual datasets per diagnostic/sex groups were excluded. Demographics and characteristics of this sample are summarized in Table 1 and in Supplementary Material in the Additional file 1.

**Table 1.**
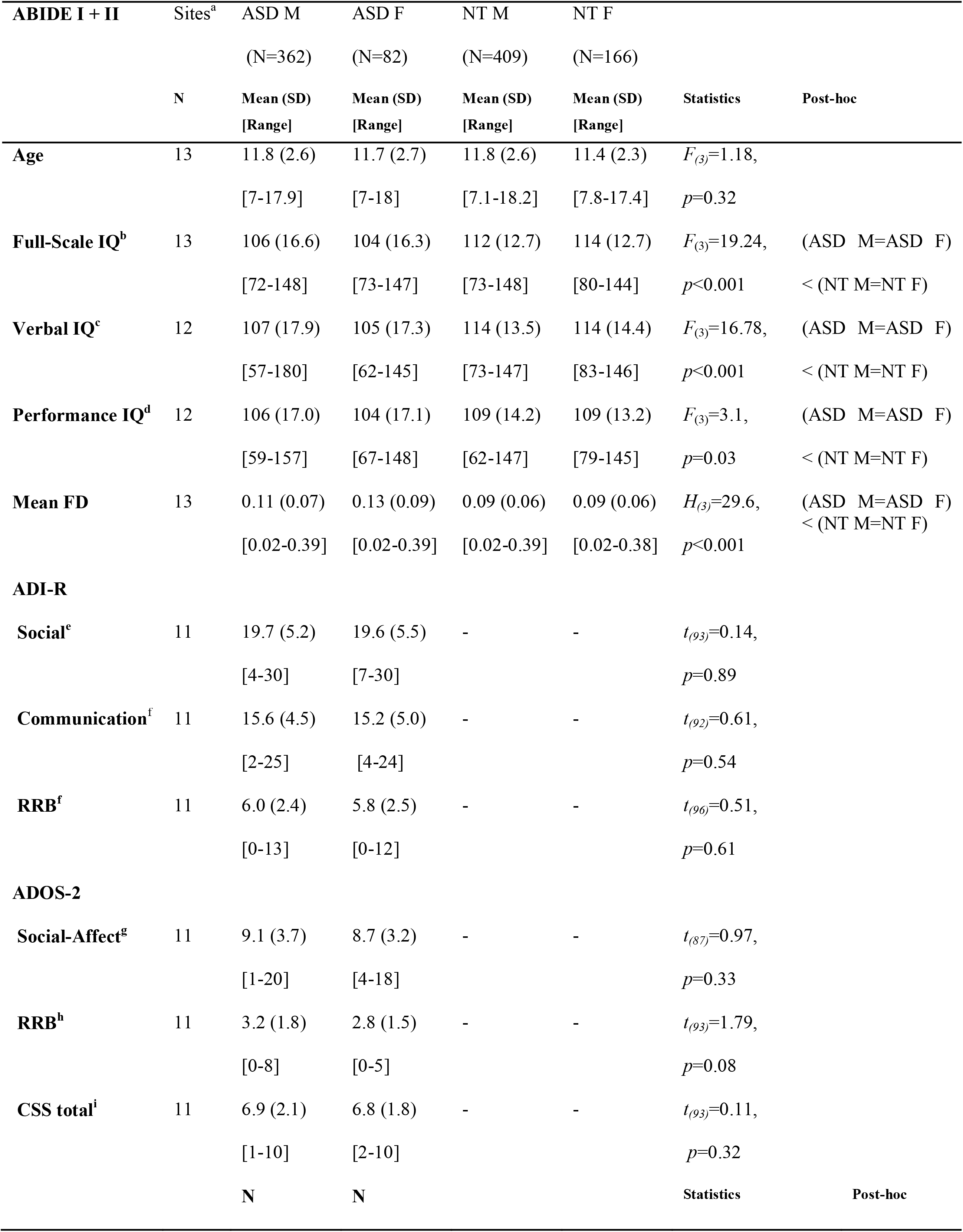
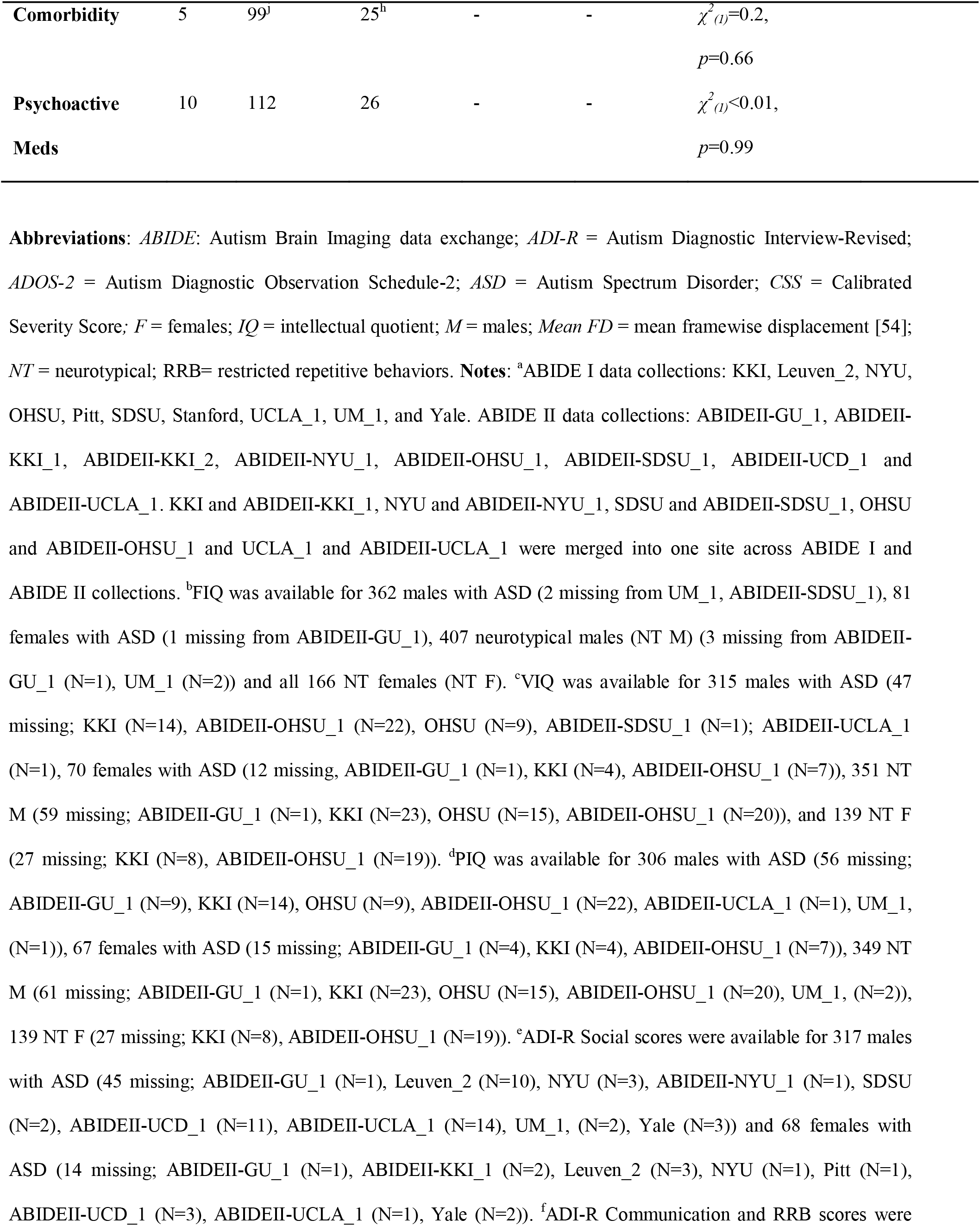
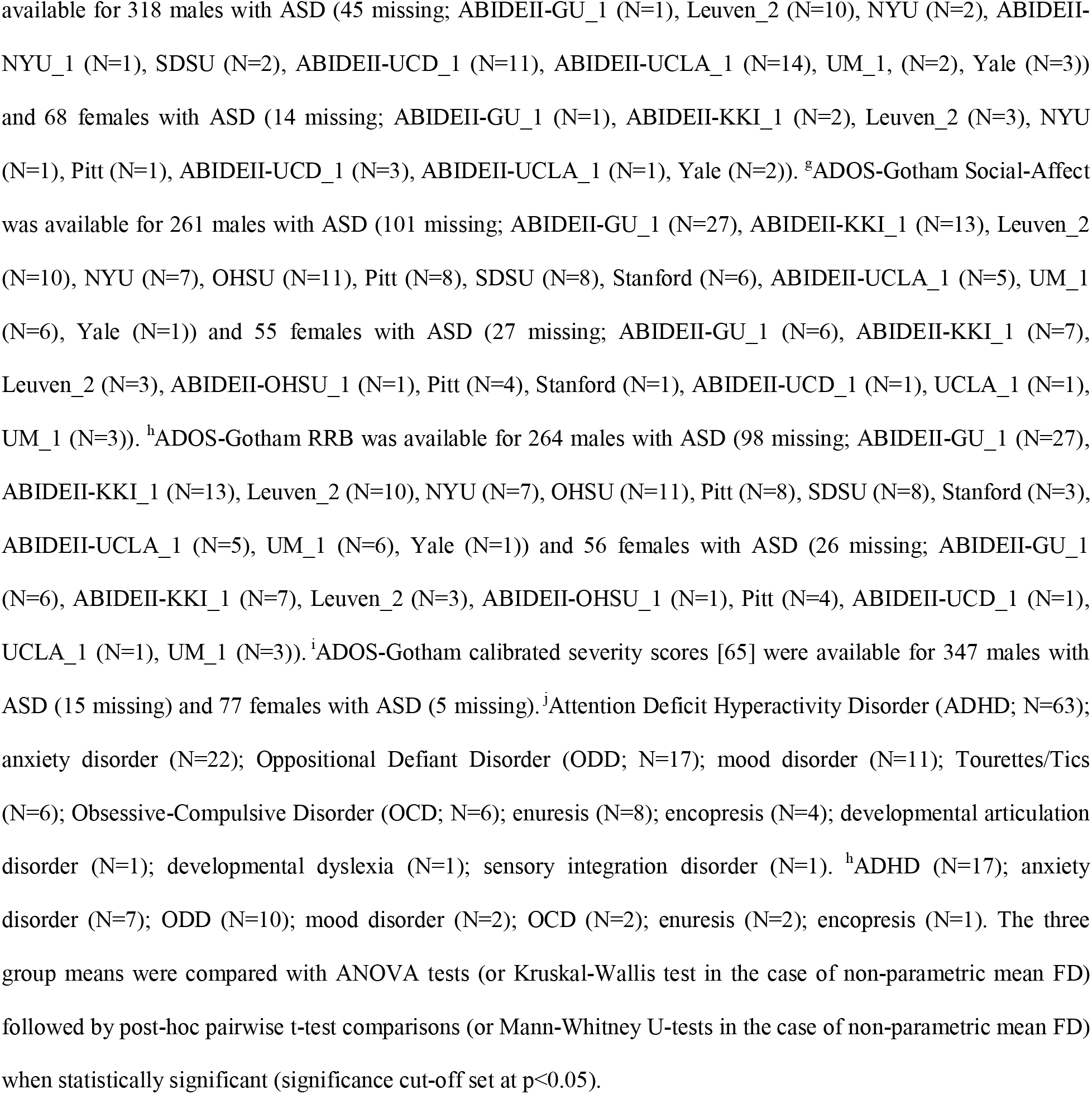
Characterization of sample merged across ABIDE I and II.

### Discovery analysis pre-processing pipeline

We examined five whole-brain R-fMRI metrics previously reported to reflect typical sex differences and found to be atypical in autism, including 1) PCC-iFC, 2) VMHC, 3) ReHo, 4) DC and 5) fALFF (see Supplementary Material, Additional file 1). R-fMRI image pre-processing steps included: slice time correction, 24 motion parameters regression [57], component-based noise reduction (CompCor) [58], removal of linear and quadratic trends, and band-pass filtering (0.01-0.1 Hz, for all metrics but fALFF). Functional-to-anatomical co-registration was achieved by Boundary Based Registration (BBR) using FSL FLIRT [59]. Linear and nonlinear spatial normalization of functional echo planar images (EPIs) to Montreal Neurological Institute 152 (MNI152) stereotactic space (2mm^3^ isotropic) was done using ANTS registration (Advanced Neuroimaging Tools) [60]. Computation of voxel-mirrored homotopic connectivity (VMHC) followed registration to a symmetric template. All R-fMRI derivatives were smoothed by a 6mm FWHM Gaussian kernel. To account for site and collection time variability across each of the data collections in ABIDE I and II data repositories, site effects were removed using the ComBat function available in python [61] (https://github.com/Jfortin1/ComBatHarmonizationhttps://github.com/brentp/combat.py). This approach has been shown to effectively account for scanner-related variance in multi-site R-fMRI data [61]. For further details see Supplementary Material in the Additional file 1.

### Discovery group-level analyses

Statistical Z-maps were generated within study-specific functional volume masks including all voxels in MNI space present across all subjects. Main effects of diagnosis and sex along with their interaction were explored by fitting a general linear model (GLM) including diagnosis or/and sex as the regressors of interest respectively, and age and mean framewise displacement (mFD) [54] as nuisance covariates. In primary analyses, we did not include FIQ as a covariate as this is thought to be suboptimal when comparing groups selected from populations carrying intrinsic IQ differences such as autism and NT [56]. Nevertheless, to provide an indication as to whether IQ may affect primary findings, in supplementary analyses FIQ was also included as an additional nuisance regressor. We applied gaussian random field theory correction based on strict voxel-level threshold of *Z*>3.1 as recommended by [62] and cluster level, *P*<0.01, given the assessment of five R-fMRI metrics in the same study (i.e., *P* 0.05/5 R-MRI metrics=0.01).

### Functional relevance of sex differences in autism

Post-hoc analyses were conducted to functionally characterize the sex-by-diagnosis interaction result(s). First, to explore the cognitive domains implicated in the cluster(s), we quantified the percentage of its overlap with 12 cognitive ontology maps [63] thresholded at *P*=1e−5. We labelled these components based on the top five tasks each component recruits [2]. Second, we used the Neurosynth Image Decoder (http://neurosynth.org/decode/) [64] to visualize the terms most strongly associated with the significant cluster. After excluding anatomical (e.g., occipital) and redundant terms (synonyms [e.g., saccades and eye movements], plurals [e.g., object and objects] or noun/adjective/adverb equivalents [e.g., vision and visual]), we visualized the top 27 terms showing correlations with the cluster map between *r*=0.64 and *r*=0.10. Third, to explore potential clinical relevance of the significant cluster, we explored brain-behavior relationships as a function of sex within the autism group. Specifically, we ran a GLM examining the interaction between biological sex and available ADOS calibrated severity total score (CSS) [65], as well as social-affect (SA) and restricted, repetitive behavior (RRB) subscores (see Supplementary Material, Additional file 1) with the dependent variable(s) being the R-fMRI metric(s) extracted from the cluster mask(s) showing a statistically significant sex-by-diagnosis effect(s).

### Robustness and Replicability

#### Robustness

We assessed whether patterns of results from the discovery analyses were observable with two other nuisance regression analytical pipelines that include commonly used data preprocessing steps. One pipeline included global signal regression (GSR) [66] which has often been used in autism studies; the other included Independent Component Analysis - Automatic Removal of Motion Artifacts (ICA-AROMA) [67] which is a relatively novel but increasingly utilized approach [46]. Given the scope of the present study, unlike prior work focusing on a wide range of individual preprocessing pipelines [46], we selected GSR and ICA-AROMA as examples of previously validated approaches thought to have impact on motion and physiological noise [45]. To assess robustness of the results observed in discovery analyses, following the voxel-level GLM, we extracted means from the masks corresponding to the same clusters that showed significant effects. These values were averaged across all the voxels in the cluster mask for a given R-fMRI metric. We used them to implement a full regression model including the predictors of interest (sex, diagnosis and their interaction), as well as age and mFD as nuisance regressors and compute effect sizes as partial eta squared (*η*_*p*_^*2*^) and their confidence intervals using the R-package ‘effectsize’. For visualization purposes we also used regressions (including sex, diagnosis, sex-by diagnosis, age, and mFD) to obtain the residuals of these mask-averaged values.

#### Replicability

Similarly, we assessed whether the group patterns observed in significant clusters identified in discovery analyses, were observed in two relatively large-scale, independent datasets selected from *a)* the EU-AIMS Longitudinal European Autism Project (LEAP), a large multi-site European initiative aimed at identifying biomarkers in autism [51,52] and *b)* the Gender Explorations of Neurogenetics and Development to Advance Autism Research (GENDAAR) dataset collected by the GENDAAR consortium and shared in the National Database for Autism Research [53]. For details on autism and NT inclusion and exclusion criteria for these samples, as well as our selection process, see Supplementary Material in the Additional file 1 [52,53]. The resulting EU-AIMS LEAP (N=309) R-fMRI datasets comprised N=133 males and N=43 females with autism as well as N=85 NT males, and N=48 NT females (see Table S1, Additional file 3); resulting GENDAAR (N=196) R-fMRI datasets comprised N=43 males and N=44 females with autism, as well as N=56 NT males and N=53 NT females (see Table S2, Additional file 3). For a comparison of demographic and clinical information between ABIDE, EU-AIMS LEAP and GENDAAR, see Table S3, S4 and S5 in the Additional file 3. After applying the same ComBat and pre-processing pipeline as used in the ABIDE-based discovery analyses, we extracted each of the R-fMRI metrics means from the Z-maps. As for robustness, we extracted values for each R-fMRI metric from the masks corresponding to the clusters showing statistically significant effects in discovery analyses and computed the corresponding effect size and residuals using the same methods described above.

For both robustness and replicability discovery findings were determined to be robust and/or replicable (R+) based on two criteria: *1)* the group mean difference(s) observed were in the same direction as those identified in the findings from discovery analyses [68] and *2)* their effects were not negligible as defined by partial eta squared *η*_*p*_^*2*^ <0.01 [69] (i.e., any small, medium or large effects) which is also consistent with prior work [41]. Finally, for consistency across analyses we also computed cluster level effect size of the discovery findings using the same approach described above.

## Results

### Discovery analyses – ABIDE

#### Main effect of diagnosis

Analyses revealed a total of seven clusters showing a significant effect of diagnosis (voxel-level *Z*>3.1; cluster-level *P*<0.01, corrected) for three of the five R-fMRI metrics: PCC-iFC (three clusters), VMHC (two clusters) and ReHo (two clusters); Figure 1, Additional file 4. These were mainly evident in anterior and posterior regions of the default network (DN) across at least two or all three R-fMRI metrics. Autism-related hypo-connectivity was present for: a) PCC-iFC, VMHC and ReHo within bilateral paracingulate cortex and frontal pole, b) VMHC and ReHo in the bilateral PCC and precuneus, and c) ReHo in right insula and central operculum (Figure 1, Additional file 4 and Additional file 5). Autism-related hyper-connectivity was only evident for PCC-iFC with left superior lateral occipital cortex, temporal occipital fusiform cortex and occipital fusiform gyrus (see Additional file 4). These results remained essentially unchanged when additionally controlling for FIQ (Additional file 6). Further, to verify that these findings were not driven by particular acquisition site(s), post-hoc analyses computed group means for diagnostic subgroups for the R-fMRI metrics extracted at the cluster-level masks excluding one out of the 13 ABIDE sites at a time. The pattern of results was essentially unchanged (Additional file 7a).

**Fig 1.**
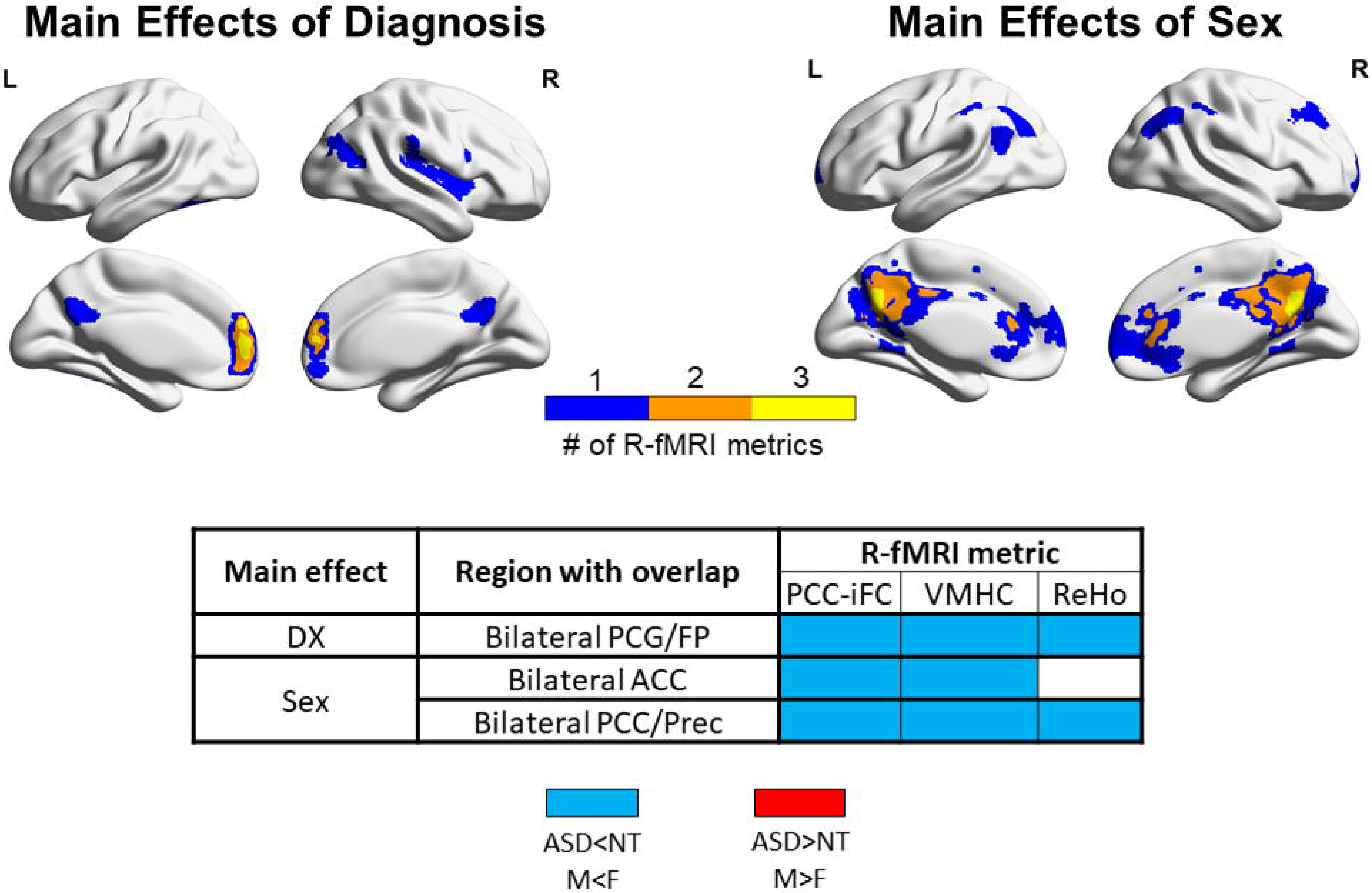
Overlap across R-fMRI metrics for main effects of diagnosis and sex. Upper panel: the surface inflated maps depict the extent of overlap across clusters showing significant main effects of diagnosis (left) and sex (right) across any of three resting state fMRI (R-fMRI) metrics showing statistically significant effects (*Z* > 3.1, *P*<0.01). Purple clusters represent areas of significant group differences emerging for only one of any of the three R-fMRI measures, orange and yellow clusters indicate measures with overlap among 2 and 3 R-fMRI measures (see Additional file 5 for statistical maps of main effects for each R-fMRI metric). Cluster masks are overlaid on inflated brain maps generated by BrainNet Viewer. Lower panel: for each of the yellow and orange clusters in panel A, the table lists the cluster’s anatomical label based on the Harvard Oxford atlas, the specific R-fMRI metrics involved, and the group difference direction (ASD<NT or M<F in blue, ASD>NT or M>F in red). Abbreviations: L= Left hemisphere; R= Right hemisphere, PCG/FP=Paracingulate cortex/frontal pole, ACC=Anterior cingulate cortex, PCC/Prec=Posterior cingulate cortex/precuneus, ASD=autism spectrum disorder, NT=neurotypical, M=Males, F=Females).

#### Main effect of sex

Analyses revealed clusters showing statistically significant main sex differences (voxel-level *Z*>3.1; cluster-level *P*<0.01, corrected), again for three R-fMRI metrics out of five in a total of 10 clusters: PCC-iFC (five clusters), VMHC (three clusters), and ReHo (two clusters). Findings involved lateral and medial portions of the DN with bilateral PCC and precuneus showing the highest overlap (see Figure 1 and Additional file 4). Specifically, regardless of diagnosis, relative to females, males showed decreased PCC-iFC with paracingulate cortex and frontal pole, right middle frontal gyrus, bilateral superior lateral occipital cortex and bilateral PCC and precuneus. Males also showed decreased VMHC and ReHo localized in PCC and precuneus. Decreased ReHo was also evident in the left angular gyrus and lateral occipital cortex in females relative to males (Figure 1, Additional file 4, and Additional file 5). These results remained essentially unchanged when additionally controlling for FIQ (see Additional file 6). Post-hoc analyses assessing the consistency of these findings across sites, as described above, revealed a similar pattern of results (Additional file 7b).

#### Sex-by-diagnosis interaction

Statistically corrected voxel-wise analyses (voxel-level *Z*>3.1; cluster-level *P*<0.01) revealed one cluster of significant sex-by-diagnosis interaction only for VMHC which was localized in the dorsolateral occipital cortex (Figure 2a and Additional file 5). Post-hoc cluster-level group means showed that NT females had higher VMHC than the three other groups, whereas autism females had lower VMHC than the three other groups (Figure 2a). Similar to the main effects, results remained essentially unchanged when additionally controlling for FIQ (Additional file 6) and analyses assessing the consistency of these findings across sites, as described above, showed a similar pattern of results (Additional file 8).

### Functional relevance of autism-related sex differences

Post-hoc analyses to functionally characterize this VMHC sex-by-diagnosis interaction indicated that the VMHC cluster in superior lateral occipital cortex overlapped with cognitive maps involved in higher-order visual, oculomotor, cognitive flexibility and language-related processes (Figure 3a). Further, as shown in Figure 3b, the most common terms were primarily related to lower-order visual processing and higher-order visual cognition, such as ‘visuospatial’ and ‘spatial attention.’ To explore potential clinical relevance of the VMHC dorsolateral occipital cluster, we explored brain-behavior relationships as a function of sex, within the autism group using three available ADOS scores (calibrated severity total score, and non-calibrated social affect and RRB subscores; see Supplementary Material in Additional file 1). Although not surviving a strict Bonferroni correction for multiple testing (i.e., 0.05/3=0.02), an interaction effect was observed for ADOS social affect scores. It revealed that more severe social deficits (*F*_*(1,311)*_=4.44, *p*=0.036) were associated with decreased VMHC in females with autism (*r*=−0.29), but not in males with autism (*r*=0.03). Given that ABIDE data were aggregated and released when calibrated social affect scores [70] were not available to assess potential differences in language abilities and age, analyses were repeated after including ADOS module (ADOS Module 2 to 4) as a nuisance covariate: results remained unchanged (*F*_*(1,306)*_=5.0, *p*=0.026) as they did also after removing the few data with the less represented ADOS module 2 (see Supplementary Material, Additional file 1). There were no significant findings, with regard to the CSS total score and non-calibrated RRB sub-score (Figure 3c), even at an exploratory statistical threshold of *P*<0.05.

**Fig 2.**
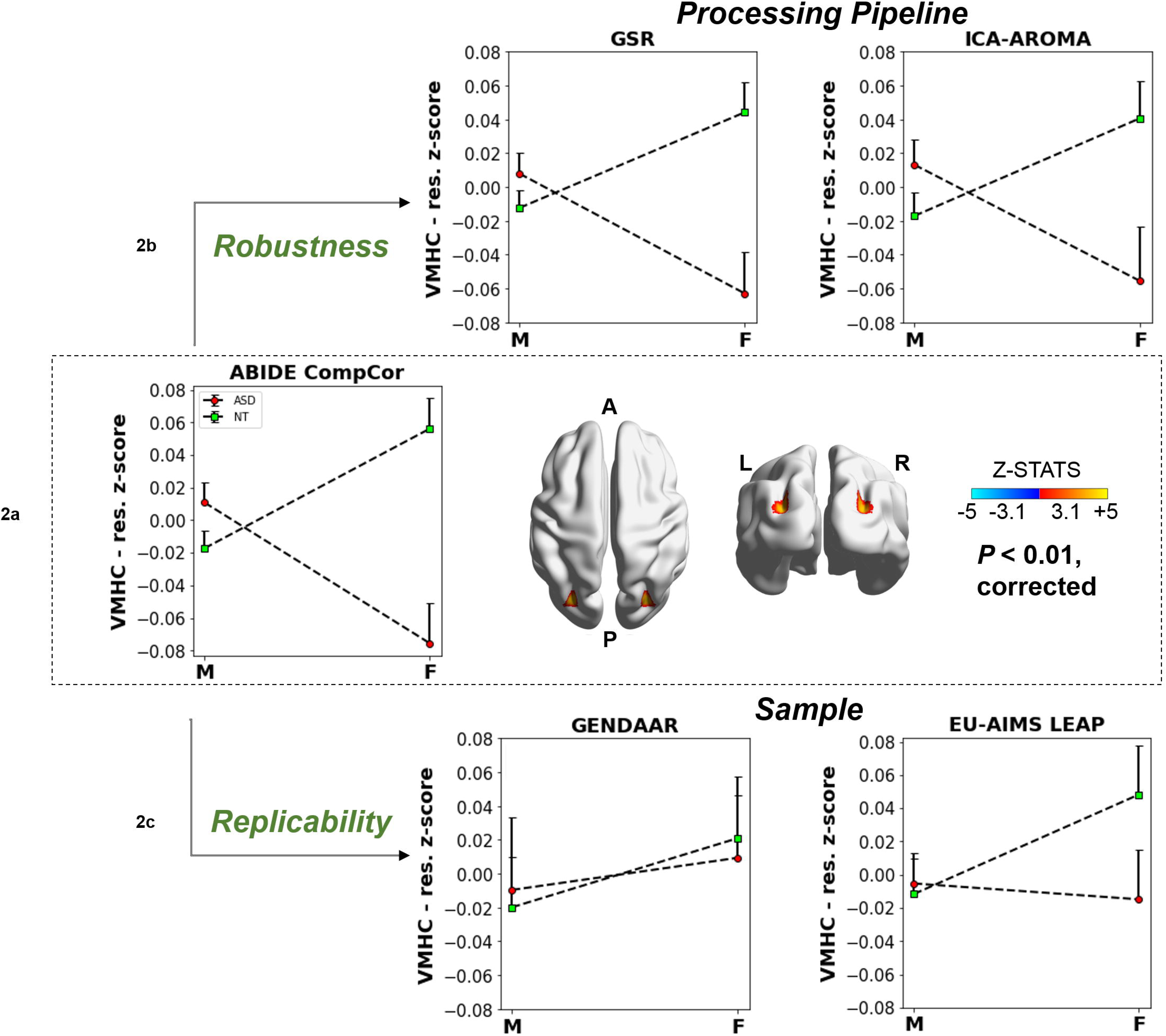
Sex-by-diagnosis interaction effect, its robustness and replicability. a) On the right, surface maps show the cluster with a significant (*Z* > 3.1, *P*<0.01) sex-by-diagnosis interaction for voxel-mirrored homotopic connectivity (VMHC) resulting from discovery analyses in the ABIDE sample using the component-based noise reduction (CompCor) pipeline. The statistical Z maps are overlaid on inflated brain maps generated by BrainNet Viewer. b) The upper panels show the pattern of VMHC group means in males and females by each diagnostic group (ASD and NT) extracted from the same cluster in data pre-processed following two alternative denoising pipelines, Global Signal Regression (GSR, left) and Independent Component Analysis – Automatic Removal of Motion Artifacts (ICA-AROMA, right). Results show a pattern similar to the those observed in discovery analyses with small to moderate effect sizes (*η*_*p*_^*2*^range=0.01–0.07). c) The lower graph shows replicability in two independent samples: the Gender Explorations of Neurogenetics and Development to Advance Autism Research (GENDAAR) and the EU-AIMS Longitudinal European Autism Project (LEAP). The pattern of results was replicable in the EU-AIMS LEAP (N=309) with a small effect size (*η*_*p*_^*2*^= 0.01) and had a negligible effect size in GENDAAR sample (N=196; *η*_*p*_^*2*^< 0.01). For all graphs VMHC data are shown as residuals obtained after regressing out mean framewise displacement and age effects. Abbreviations: L=left, R=right, A=anterior, P=posterior.

**Fig 3.**
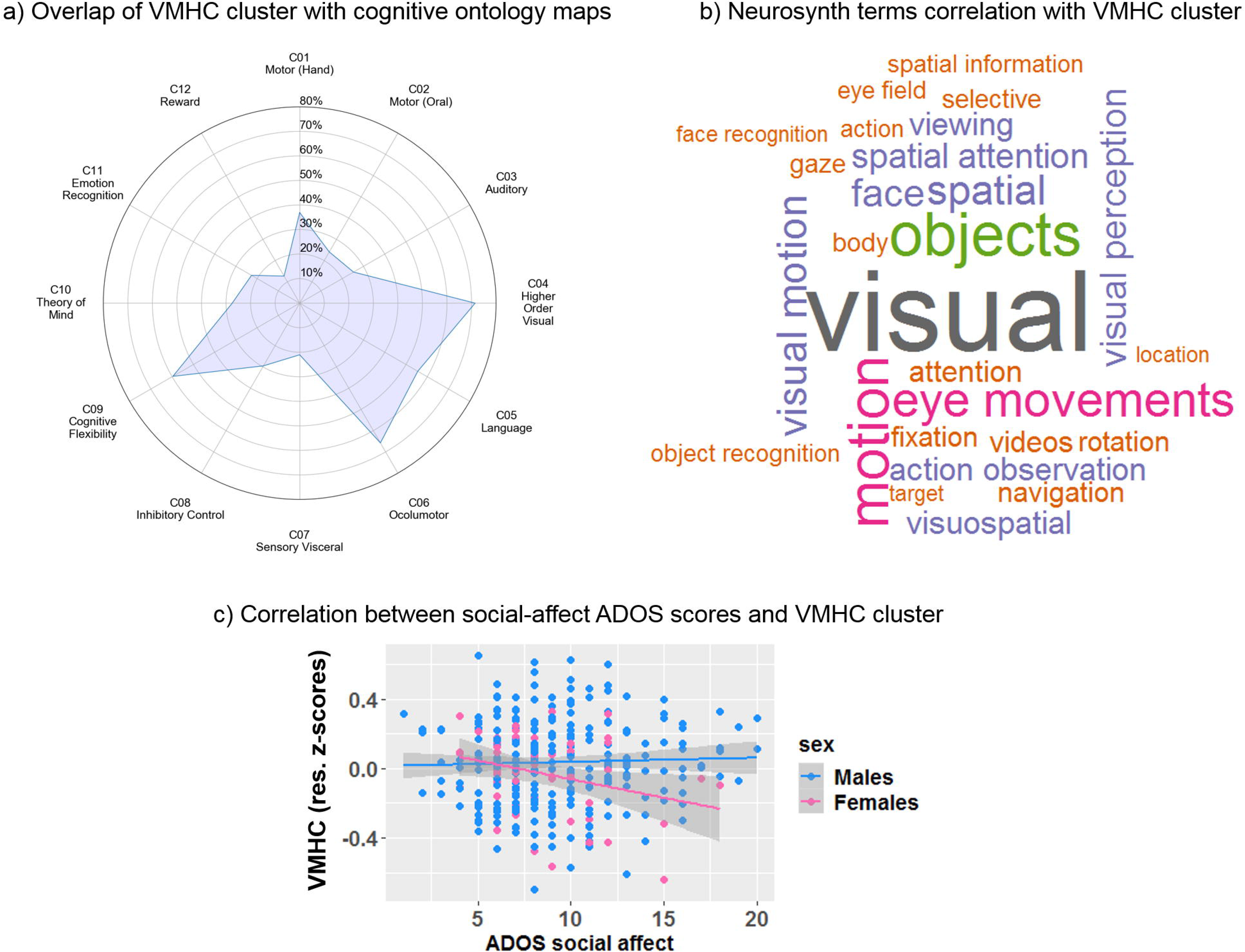
Functional relevance of sex-by-diagnosis interaction in VMHC. a) The radar plot shows the percentage (0-80%) of overlap between the voxels in the dorsolateral occipital cluster showing a significant VMHC sex-by-diagnosis interaction in discovery analyses and the 12 Yeo cognitive ontology probability maps [63] (probability threshold at *P* = 1e-5) for cognitive components C1-C12. As in Floris et al. [2], we labelled each component based on the top five tasks reported to be most likely recruited by a given component. b) Word cloud based on the top 27 terms showing correlations between *r*=0.64 to *r*=0.10 associated with the same VMHC cluster based on the Neurosynth Image Decoder. c) Sex-differential association between each individual’s VMHC at the cluster showing a significant sex-by-diagnosis interaction in primary analyses and available ADOS social-affect uncalibrated sub-scores in males and females with ASD. VMHC data are shown as residuals obtained after regressing out mean framewise displacement and age effects. While males showed no significant associations at corrected and uncorrected thresholds, females with lower dorsolateral occipital VMHC showed more severe social-affect symptoms at uncorrected statistical threshold (*F*_*(1,311)*_=4.44, *p*=0.036).

### Robustness

The same pattern of results identified in discovery analyses was observed in the results pre-processed using GSR or ICA-AROMA, across the three R-fMRI metrics in all the clusters identified in the primary analyses across main effects of diagnosis, sex and their interaction; effect size ranges from small to moderate as in discovery analyses (*η*_*p*_^*2*^ range=0.01–0.07; Figure 2b, Figure 4, Additional file 5, Additional file 9).

**Fig 4.**
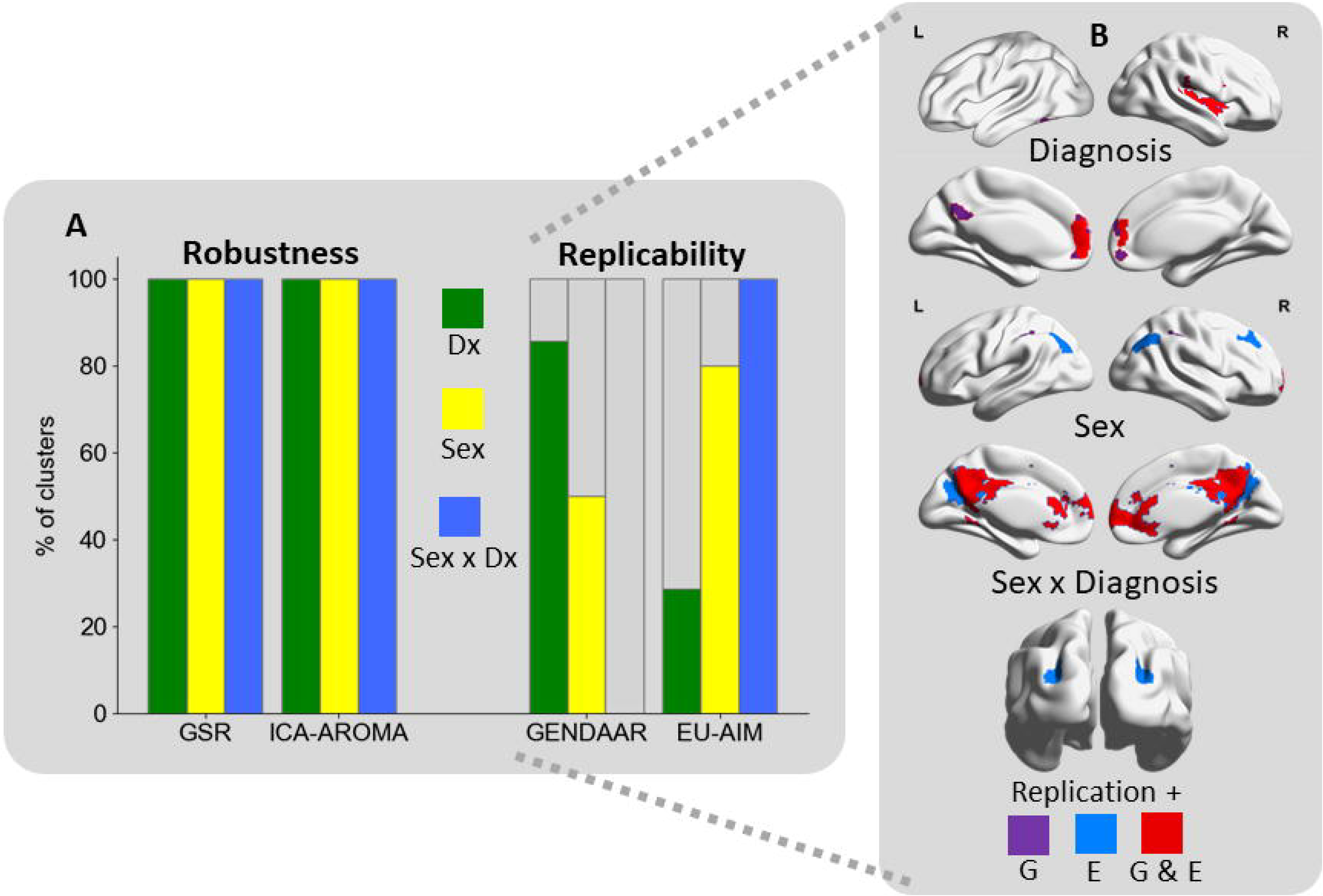
Robustness and replicability summary. a) The histogram summarizes the percentage of clusters showing a robust and replicable pattern of results as that observed in discovery analyses in the ABIDE sample for main effects of diagnosis (Dx; green; N=7 clusters), sex (yellow; N=10 clusters) and their interaction (blue; N=1 cluster) across three R-fMRI metrics. All findings were robust to different preprocessing pipelines. Across R-fMRI metrics, main sex effects were moderately (50%) to largely (80%) replicable across independent samples: Gender Explorations of Neurogenetics and Development to Advance Autism Research (GENDAAR) and the EU-AIMS Longitudinal European Autism Project (LEAP), respectively. Replicability for main effects of diagnosis was largely replicable in GENDAAR (86%) and minimally replicable in EU-AIMS LEAP (29%). The VMHC pattern observed for sex-by-diagnosis interaction in primary discovery analyses was replicated in EU-AIMS LEAP only. b) Surface conjunction maps show the clusters replicated in GENDAAR only (G, purple), EU-AIMS LEAP only (E, blue) and in both samples (G&E, red) for each effect separately. Cluster masks are overlaid on inflated brain maps generated by BrainNet Viewer.

### Replicability

#### Main effects of diagnosis

Main effects of diagnosis showed higher replicability (i.e., non-negligible *η*_*p*_^*2*^ effects showing a similar group mean pattern as observed in discovery analyses) in GENDAAR than in EU-AIMS LEAP. Specifically, across the three R-fMRI metrics that showed significant diagnostic differences in discovery analyses, six of the seven clusters (86%) in GENDAAR, were replicated (*η*_*p*_^*2*^ range=0.01–0.04); only two of those seven (29%) were replicated in EU-AIMS LEAP (*η*_*p*_^*2*^ range=0.01–0.04). Nevertheless, clusters showing decreased ReHo in ASD vs. NT across the insula and central operculum, as well as in the frontal pole were replicated across all samples (Figure 4, Additional file 5, Additional file 10a).

#### Main effects of sex

Across all three R-fMRI metrics, the main effects of sex observed in primary discovery analyses was evident in both independent samples, for most clusters in the EU-AIMS LEAP (80%; 8/10) with effects size ranging from small to moderate (*η*_*p*_^*2*^ range=0.01–0.06) and for half of the clusters in GENDAAR (50%; 5/10), albeit with small effects (*η*_*p*_^*2*^ range=0.01–0.02); Figure 4, Additional file 5, Additional file 10b). Notably, the pattern of typical sex differences localized along the default network midline (i.e., decreased VMHC, ReHo and PCC-iFC) was replicated across both independent samples (Figure 4b).

#### Sex-by-diagnosis interaction effect

The pattern of autism-related VMHC sex differences observed in discovery analyses in the superior lateral occipital cortex was observed in the EU-AIMS LEAP dataset, (*η*_*p*_^*2*^ =0.01); (Figure 2c, Figure 4, Additional file 5). In the GENDAAR dataset, while group means in males with autism, NT males and NT females showed a similar direction as in the ABIDE discovery findings, females with autism differed in magnitude and in the direction of group differences, resulting in a negligible effect with *η*_*p*_^*2*^ <0.01. (Figure 2c).

## Discussion

We examined autism-related sex differences for intrinsic functional brain organization across multiple R-fMRI metrics in a large discovery sample of males and females with autism relative to age-group matched NT selected from the ABIDE repositories [22,23]. Analyses revealed significant main effects of sex and diagnosis across intrinsic functional connectivity (iFC) of the posterior cingulate cortex, regional homogeneity and voxel-mirrored homotopic connectivity (VMHC) in several cortical regions. Notably, main effects converged along the midline of the default network. In contrast, sex-by-diagnosis interactions were limited to VMHC in the superior lateral occipital cortex. Placed in the context of sex and diagnostic main effects on interhemispheric homotopic connectivity in cortical regions, this result, suggests that atypical interhemispheric interactions are pervasive in autism but reflect a combination of sex-independent (i.e., main effect of diagnosis common across both sexes) and sex-dependent (i.e., sex-by-diagnosis interaction) effects, each specific to different functional cortical system. This sex-by-diagnosis interaction effect was robust to distinct pre-processing strategies as those observed for main effects. Further, despite the lack of *a priori* harmonization for data acquisition among the three samples, this finding was replicable in the larger of the two independent samples (i.e., EU-AIMS LEAP). On the one hand, this, together with largely replicable main effects of sex with variable replicability of main diagnostic effects by sample, suggests that inter-sample replicability of R-fMRI can be feasible in autism when sources of variability in diagnostic groups are accounted for in samples sized properly to address such variability. On the other hand, our results highlight the urgent need to obtain multiple harmonized datasets properly powered to systematically address and understand sources of heterogeneity, including and beyond the role of biological sex.

### Sex-dependent and independent atypical interhemispheric interactions in autism

VMHC reflects inter-hemispheric homotopic relations. Its strength has been suggested to index coordinated cross-hemispheric processing: *stronger* VMHC index weaker hemispheric specialization and vice versa [33,71]. Several lines of evidence support the notion that the neurobiology of autism is related to atypical hemispheric interactions, including homotopic connectivity and hemispheric lateralization [35,72,81,73–80]. VMHC and functional hemispheric lateralization have also been shown to be sex-differential in NT [33,82,83]. The dorsolateral occipital association cortex identified in our discovery analyses is known to serve hemispherically specialized processes, such as visuospatial coordination [84]. Thus, our findings of NT males’ VMHC in dorsolateral lateral occipital cortex being lower than NT females are consistent with the notion of increased hemispheric lateralization in this cortical region in NT males relative to NT females. In our data, females with autism instead showed even lower VMHC than NT males, while males with autism showed slightly higher VMHC than NT males. This pattern is indicative of ‘gender-incoherence’ [20] as males and females with autism display the opposite pattern expected in NT per their biological sex. Findings of ‘gender incoherence’ have been reported in earlier neuroimaging studies of autism using different modalities [3,85,86]. Among them, several R-fMRI studies explicitly focusing on detecting diagnosis-by-sex interactions (i.e., the regression model included a sex-by-diagnosis interaction term) [3,9] yielded a pattern of results consistent with ‘gender incoherence.’ In contrast, other studies [10–12] reported a pattern consistent with the ‘extreme male brain theory’ [19] – i.e., a shift towards maleness in both females and males with autism. While the seemingly diverging conclusions of these two sets of studies may be attributed to methodological differences, such as the extent of brain networks explored and the statistical modelling employed, findings from our prior work suggest that both shifts towards either maleness or femaleness co-occur in the intrinsic brain of males with autism, in a network-specific manner [2]. However, such prior work did not include female data. Thus, by not directly assessing sex-by-diagnosis interactions, unlike the present study, results could not point to patterns affecting diagnostic differences between the sexes vs. those that are common to autism across both sexes [4]. This is relevant for efforts focusing on identifying underlying mechanisms. Findings resulting from sex-by-diagnosis interactions may shed light on sex differential mechanisms that go awry in autism and may reflect sex-specific susceptibility mechanisms. On the other hand, diagnostic atypicalities common for both sexes may reflect factors central to the emergence of autism, regardless of whether they overlap with patterns known to be differential between sexes [87]. Interestingly, a recent study based on a sample selected from GENDAAR [14] revealed that the iFC between the nucleus accumbens (selected *a priori*) and a region of the dorsolateral occipital cortex partially overlapping with that identified by our VMHC analyses, was differentially modulated by the aggregate number of oxytocin receptor risk alleles in females with autism vs. NT females and vs. males with autism. Although VMHC was not directly tested in that earlier study [14], its result in dorsolateral occipital cortex is consistent with our observation of atypical sexual differentiation of this visual network region and, together, suggest the need for future whole-brain studies of oxytocin effects in autism.

Along with the sex-dependent autism patterns, our analyses found statistically significant main effects of diagnosis in inter-hemispheric interactions indexed by VMHC in distinct cortical circuits. These were localized along the midline of the DN (paracingulate/frontal cortex consistently and PCC/precuneus) where main effects of PCC-iFC and ReHo also converged. Our results are consistent with prior reports of atypical intrinsic organization of the DN in autism [10,23,26,88–90]. Together they support the role for a common, sex-independent DN role in autism. This is also supported by a recent autism neurosubtyping study that identified three latent iFC factors, all sharing DN atypicalities along with their neurosubtype-specific patterns [91]. Building on this evidence to disentangle the specific role of each of the factors affecting autism in sex-independent and sex-dependent ways, a necessary next step is to engage in novel large-scale data collection efforts including more female data.

### Robustness, replicability and sources of variability

The growing awareness of the replication crisis in neuroscience [92–94] motivated our analyses examining robustness and replicability of findings. While a comprehensive and systematic reproducibility assessment is beyond the scope of the study, here we focused on examining whether the findings observed in the discovery analyses were also seen after using different preprocessing pipelines – *robustness* – as well as in fully independent, albeit of convenience samples (i.e., not harmonized *a priori* with each other) – *replicability*. To this end, given the lack of consensus on quantitative metrics of replicability, we opted to use measures of effect size. These are considered complementary to null hypothesis significance testing [95]. In the context of this study, given the use of convenience samples of different sizes, their selection was considered an advantageous and practical means to provide information on the magnitude of group differences in diagnosis and sex, as well as their interaction. Here, we considered findings to be robust and/or replicable for any non-negligible effects (i.e., *η*_*p*_^*2*^≥ 0.01; [69]). We reasoned that given their distributed and heterogenous nature [6], atypicalities in the autism connectome can stem from a combination of differently sized non-negligible effects, as shown for autism in other biological domains such as genetics [96,97].

With this in mind, across the two preprocessing methods examined here, the patterns of findings were consistent with those observed in discovery analyses across all R-fMRI metrics and effects. These robustness results are consistent with a prior study by He and colleagues [46] reporting that differences in a wider range of pre-processing pipelines have marginal effects on variation in diagnostic group average comparisons. Our study confirms and builds on this earlier report by extending findings of robustness to sex group mean differences and their interactions with diagnosis.

A more nuanced picture emerged from the inter-sample analyses as replicability varied by sample, across the effects and R-fMRI metrics examined. Specifically, while inter-sample main effects of sex were moderately to largely replicable across R-fMRI metrics on both independent samples (~50 to 80% of the clusters in GENDAAR and EU-AIMS-LEAP, respectively), replicability of diagnostic effects significantly varied by sample (86% to 29%) across R-FMRI metrics. This is at least in part consistent with findings by King et al. [50] also showing that, depending on the R-fMRI feature examined, diagnostic group differences varied across samples. Even in this scenario, King et al. also reported that findings of decreased homotopic connectivity in autism were relatively more stable than other R-fMRI metrics [50]. This observation, combined with replicability of our VMHC sex-by-diagnosis interaction findings in the larger of the two independent samples (EU-AIMS LEAP), suggests that measures of homotopic connectivity may have a relevant biological relevance for autism. It is also possible that given its moderate to high test-retest reliability, VMHC is more suitable in efforts assessing replicability [98,99].

The striking clinical and biological heterogeneity in autism should be considered as a major contributor to discrepancies in findings of studies focusing on the main effects of diagnostic group means contrasts/interactions [100–103]. Against this background, we interpret our replicability findings on diagnostic effects and, in turn, diagnosis-by-sex interactions. Inter-sample differences may have contributed to the more variable results of replicability on the diagnosis main effects. These may include autism symptom level, age, and IQ, albeit secondary analyses suggested that the examined IQ range did not substantially affect the pattern of discovery results. For example, the EU-AIMS LEAP sample was on-average older, had lower VIQ and most notably, lower symptom severity across all subscales of the ADOS and ADI-R than the ABIDE sample. On the other hand, the GENDAAR sample (which has greater number of replicable diagnostic mean group patterns) did not differ from ABIDE in these variables, except for mean age. Furthermore, a fact that is often neglected, is that the NT groups may also present with considerable sample heterogeneity between studies [100,104]. For instance, our NT controls in the EU-AIMS LEAP sample had lower VIQ than both ABIDE and GENDAAR NT controls. This has potentially influenced the low replicability of diagnosis main effects in EU-AIMS LEAP.

In contrast, sex-by-diagnosis effect on VMHC in the dorsolateral occipital cortex was replicable in the larger samples, the EU-AIMS LEAP, but not in GENDAAR. Small samples introduce larger epistemic variability (i.e., greater variation related to known and unknown confounds) [105]. Increasing the number of subjects/data allows mitigating epistemic variability and, thus, capturing the underlying variability of interest. Thus, although the rate of EU-AIMS LEAP replicability for the main effect of diagnosis was limited, accounting for biological sex, a known key source variability in autism, may have contributed to a replicable sex-by-diagnosis pattern in this larger sample. In line with sample size concerns, using four datasets sized between 36 and 44 individuals selected from the ABIDE repository, He et al. [46] found low similarity rates of diagnostic group-level differences on the strength of iFC edges in contrast with the largely similar pattern of results across pipelines. Of note, unlike prior efforts [46,50], we controlled for site effects within each of the samples (i.e., ABIDE, GENDAAR and EU-AIMS LEAP), using ComBat. Future large-scale harmonized data collections are needed to control and assess the impact of inter-sample variability. Taken together, these findings highlight that sample differences can impact replicability.

Beyond clinical and biological sources of variation, samples may differ in MRI acquisition methods, as well as in approaches used to mitigate head motion during data collection and its impact on findings [106]. Adequately controlling for head motion remains a key challenge for future studies assessing inter-sample replicability. For the present study, we excluded individuals with high motion, retained relatively large samples with group average low motion (mean α standard deviation range mFD range=0.09-0.16 α 0.06-0.10 mm), as well as included mFD at the second-level analyses as a nuisance covariate. Overall, the extent to which each sample-related factor affects replicability needs to be systematically examined in future well-powered studies. Only this type of studies will allow for emerging subtyping approaches to dissect heterogeneity by brain imaging features using a range of data-driven methods [107,108], including normative modelling [109–111].

Inter-sample differences and methodological differences, beyond nuisance regression, may have contributed to some differences in findings between the present and earlier studies, conducted with independent or partially overlapping samples [9,23,41]. For example, Alaerts et al. [9] also examined sex-by-diagnosis interaction in PCC-iFC in a dataset selected from ABIDE I only. Although their pattern of results was consistent with the ‘gender incoherence’ model, the resulting circuit(s) did not involve the dorsolateral occipital cortex as identified with VMHC in the present study. Along with differences in samples selected from the same data repositories, other ABIDE-based methodological choices that may affect results. For example, prior studies differed with the present one in the inclusion of sex-by-diagnosis interaction [15], the extent of the whole-brain voxel-based analyses [13,14], or the statistical threshold utilized [23]. Nevertheless, it is remarkable that even in light of these differences, consistent results have emerged including the role of the ‘gender incoherence’ model for females with autism, atypical inter-hemispheric interactions in autism, and sex-dependent and sex-independent atypical intrinsic brain function across distinct functional networks.

### Limitations

Along with the inter-sample differences resulting from the lack of sufficiently available harmonized inter-site replication datasets in the field, other limitations of this study should be addressed in future efforts. One regards the lack of measures differentiating the effects of sex vs. that of gender so as to disentangle their relative roles (e.g., gender-identity and gender-expression) in the intrinsic brain properties [112]. Further, in-depth cognitive measures to directly characterize the role of VMHC findings were not available. Additional behavioral measures are needed to establish whether our result in VMHC of the dorsolateral occipital cortex mainly applies to low-level (bottom-up) visual processing differences or higher-level (top-down) attentional/controlled processes in males and females with autism. As a neurodevelopmental disorder, autism shows striking inter-individual differences in clinical and developmental trajectories, as well as outcomes. Thus, age may influence symptom presentation [113,114] and neurobiology [11,110,115]. Despite the considerable size of the samples available for this study, it is still difficult to sufficiently cover a broad age range across both males and females and diagnostic groups across contributing sites and to evaluate age effects appropriately. Even larger cross-sectional samples are needed to derive meaningful age-related information that ultimately require confirmation in longitudinal study designs. Such longitudinal studies would allow to examine the potential impact of puberty and related surge of sex steroids reported in NT boys and girls [116], in autism specifically. Here, we have no direct measure of puberty other than age, but future studies should aim to include such measures. Further, the value of a large-scale and publicly available multi-site resource such as ABIDE also comes with unavoidable differences in site differences which must be considered in data selection, analyses and interpretation of results. Although residual site-related effects may have remained in findings even after using the novel Bayesian approach for correcting for batch-effects, replicability in independent samples suggest that effects are not simply driven by site variability. These results are consistent with earlier reports of reproducible imaging biomarkers even when accounting for inter-site differences in multisite datasets such as the ABIDE I repository [47]. Finally, despite the advantages of effect sizes over p-value when comparing independently collected samples of different sizes and potentially different variances, it is important to acknowledge they are not without limitations [117] and should be interpreted with caution. Similar to p-values, they are dependent on sample sizes and have the equivalent risks of p-hacking. Finally, effect sizes standard errors can be large - a concern we addressed through including of confidence intervals.

## Conclusions

The present work revealed sex differences in the intrinsic brain of autism, particularly in dorsolateral occipital interhemispheric interactions, which were robust to pre-processing pipeline decisions and replicable in the larger of the two independent samples. While differences in nuisance regression pipelines have little influence on the consistency of findings, sample heterogeneity represents a challenge for replicability of findings. Lateralized cognitive functions and cross-hemispheric interactions should be further explored in relation to sex differences in autism while addressing this challenge with future harmonized data acquisition efforts with even larger samples.

## List of abbreviations

ABIDE: Autism Brain Imaging Data Exchange
ASD: Autism spectrum disorder
DC: Degree centrality
DN: Default network
EPI: Echo-planar image
LEAP: Longitudinal European Autism Project
fALFF: Fractional amplitude of low frequency fluctuations
FIQ: Full-scale IQ
GENDAAR: Gender Explorations of Neurogenetics and Development to Advance Autism Research
GSR: Global Signal Regression
ICA-AROMA: Independent Component Analysis - Automatic Removal of Motion Artifacts
iFC: Intrinsic functional connectivity
mFD: Mean framewise displacement
NDAR: National Database for Autism Research
NT: Neurotypical
PCC: Posterior cingulate cortex
ReHo: Regional Homogeneity
R-fMRI: Resting-state functional magnetic resonance imaging
VMHC: Voxel-mirrored homotopic connectivity

## Ethics approval and consent to participate

ABIDE I and II: All contributions were based on studies approved by the local Institutional Review Boards, and data were fully anonymized (removing all 18 HIPAA (Health Insurance Portability and Accountability)-protected health information identifiers, and face information from structural images).

EU-AIMS LEAP: Ethical approval was obtained through ethics committees at each site. GENDAAR: Informed assent and consent were obtained from all participants and their legal guardians, and the experimental protocol was approved by the Institutional Review Board at each participating site.

## Consent for publication

Not applicable

## Availability of data and materials

*ABIDE I and II*: Data are freely accessible in the publicly available Autism Brain Imaging Data Exchange repository (http://fcon_1000.projects.nitrc.org/indi/abide).

*EU-AIMS LEAP*: Data are currently only available for sites involved in data collection, but data (starting with the first wave) will be made available on request.

*GENDAAR*: Anonymized data are publicly available through the National Database for Autism Research (NDAR) and access can be requested via https://nda.nih.gov/about.html.

## Authors’ contributions (in alphabetical order by contribution)

ADM and DLF have contributed to the study’ s conception; ADM, DLF, JOAF, M-CL, MPM have contributed to distinct aspects of the study design and interpretation of all findings; DLF and JOAF conducted the analyses; SG has provided support in all data analyses requiring CPAC; ADM, DLF, JOAF have generated figures and tables; ADM and DLF drafted the manuscript; ADM, DLF, JOAF, M-CL, and MPM have revised and edited multiple versions of the manuscript; CFB, MM, MO, have organized the EU-AIMS LEAP data and edited latest manuscript versions and its revisions; CFB, CE, CM, DGMM, EL, FDA, GD, JKB, JT, SB-C, TC, TB, SD have contributed to the coordination, data acquisition and coordination of the EU-AIMS LEAP project, as well as edited later versions of the manuscript and its revisions. All authors read and approved the manuscript.

## Acknowledgements

The authors thank all investigators and contributors to the Gender Explorations of Neurogenetics and Development to Advance Autism Research (GENDAAR) for collecting and sharing their data as well as addressing question related to the data, the Autism Brain Imaging Data Exchange, and the contributors to EU-AIMS Longitudinal European Autism Project for their efforts in data collection and sharing. The GENDAAR Consortium comprises, in alphabetical order, Elizabeth H. Aylward, Raphael A. Bernier, Susan Y. Bookheimer, Mirella Dapretto, Nadine Gaab, Daniel H. Geschwind, Andrei Irimia, Allison Jack, Charles A. Nelson, Kevin A. Pelphrey, Matthew W. State, John D. Van Horn, Pamela Ventola, and Sara J. Webb. We thank all participants and their families for participating in the respective study.

## Additional files

### Additional file 1

**Title:** Supplementary Material

**File format:** docx

**Description:** Supplementary Methods

### Additional file 2

**Title:** Selection flowchart for the ABIDE sample

**File format:** tif

**Description:** The flowchart illustrates the selection process resulting in the final ABIDE I and II combined sample of 1019 subjects. At each flowchart step, the numbers outside the parentheses represent the total number of datasets across both ABIDE I and ABIDE II; in parenthesis are the number of datasets derived from ABIDE I (the resulting difference between these numbers would be the numbers for dataset stemming from ABIDE II). The rationale for each selection step is detailed in Supplementary Material. Abbreviations: ASD=autism spectrum disorder, NT=neurotypical, A I=ABIDE I.

### Additional file 3

**Title:** Supplementary Tables

**File format:** docx

**Description:** Characterization of EU-AIMS LEAP and GENDAAR samples, comparison between samples and summary table of main effects.

### Additional file 4

**Title:** Main effects of diagnosis and sex in the ABIDE discovery sample

**File format:** tif

**Description:** Significant results (*Z* >3.1, *P*<0.01, corrected) of voxel-wise discovery analyses conducted in the ABIDE dataset for main effects (ME) of diagnosis (left) and sex (right) for seed-based intrinsic functional connectivity of the posterior cingulate cortex- (PCC), voxel mirror homotopic connectivity (VMHC), and Regional Homogeneity (ReHo). Significant clusters are overlaid on inflated brain maps generated by BrainNet Viewer. No significant effects were detected for degree centrality or fractional amplitude of low frequency fluctuations. ME Diagnosis: <u>PCC-iFC:</u> bilateral paracingulate cortex and frontal pole (PCG/FP), superior lateral occipital cortex (sLOC), temporal occipital fusiform cortex and occipital fusiform gyrus (TOFC/OFC); <u>VMHC</u>: bilateral posterior cingulate gyrus and precuneus (PCC/Prec), PCG/FP; <u>ReHo</u>: PCG/FP, central operculum and insula (CO/Ins). ME Sex: <u>PCC-iFC</u>: bilateral sLOC, middle frontal gyrus (MFG), bilateral PCC/Prec, bilateral PCG/FP; <u>VMHC</u>: bilateral PCC/Prec, bilateral anterior cingulate cortex (ACC); <u>ReHo</u>: bilateral PCC, angular gyrus and lateral occipital cortex (AnG/LOC). See Additional file 3: Table S6 for details on each cluster sizes. *Due to processing failure of two subjects for VMHC, the sample size comprised 1017 subjects instead of 1019.

### Additional file 5

**Title:** Characteristics of the clusters with significant effect and effect size across analyses

**File format:** pdf

**Description:** Additional file 5 summarizes cluster’ anatomical labels, center of gravity coordinates and statistics derived from discovery analyses, as well as effect size and their confidence of interval for each of these clusters across all analyses In green are the effect found to be robust/replicable based on our criteria (*i.e.,* the group mean difference(s) in the same direction as those identified in discovery analyses and effect sizes not negligible as defined by partial eta squared *η*_*p*_^*2*^ <0.01) and in yellow those that did not. PCC-iFC: posterior cingulate cortex intrinsic functional connectivity, VMHC: voxel-mirrored homotopic connectivity, ReHo: regional homogeneity, TOFC/OFG: temporal occiptal fusiform cortex/occiptal fusiform gyrus, sLOC: superior lateral occipital cortex, PCG/FP: paracingulate cortex/frontal pole, PCC/Prec: posterior cingulate gyrus/precuneus, CO/Ins: central operculum/insula, MFG: middle frontal gyrus, ACC: anterior cingulate cortex, SMG: supramarginal gyrus, AnG/LOC: angular gyrus/lateral occipital cortex. *Due to processing failure of two subjects for VMHC, the sample size comprised 1017 subjects instead of 1019 for ABIDE and 307 instead of 309 for EU-AIMS.

### Additional file 6

**Title:** Main effects of diagnosis and sex in the ABIDE discovery sample when additionally covarying for FIQ

**File format:** tif

**Description:** Including full-scale IQ (FIQ) as a nuisance regressor in addition to age and mean FD in the voxel-wise model yielded significant (*Z*>3.1, *P*<0.01, corrected) findings highly similar to those observed in discovery analyses across main effects (ME) of diagnosis (left) and sex (right), sex and their interaction. As in the discovery approach, analyses were conducted for seed-based intrinsic functional connectivity of the posterior cingulate cortex-(iFC-PCC), voxel mirror homotopic connectivity (VMHC), and Regional Homogeneity (ReHo). Significant clusters are overlaid on inflated brain maps generated by BrainNet Viewer. No significant effects were detected for degree centrality or fractional amplitude of low frequency fluctuations. ME Diagnosis: <u>PCC-iFC:</u> bilateral paracingulate cortex and frontal pole (PCG/FP), superior lateral occipital cortex (sLOC), temporal occipital fusiform cortex and occipital fusiform gyrus (TOFC/OFC); <u>VMHC</u>: bilateral posterior cingulate gyrus and precuneus (PCC/Prec), PCG/FP; <u>ReHo</u>: PCG/FP, central operculum and insula (CO/Ins). ME Sex: <u>PCC-iFC</u>: bilateral sLOC, middle frontal gyrus (MFG), bilateral PCC/Prec, bilateral PCG/FP; <u>VMHC</u>: bilateral PCC/Prec, bilateral anterior cingulate cortex (ACC); <u>ReHo</u>: bilateral PCC, angular gyrus and lateral occipital cortex (AnG/LOC). Sex-by-diagnosis: VMHC: bilateral dorsolateral occipital cortex. *Due to processing failure of two subjects for VMHC, the sample size comprised 1017 subjects instead of 1019.

### Additional file 7

**Title:** Stability of main effects

**File format:** tif

**Description:** Inter-site stability was assessed after extracting group means at masks corresponding to the clusters showing significant main effects of diagnosis (7a) and sex (7b) in the discovery analyses and then deriving the group mean when leaving one acquisition site out at the time. The pattern of results was unchanged. Different ABIDE sites are color-coded on legend on the side. Due to processing failure of two subjects for VMHC, the sample size comprised 1017 subjects. Abbreviations: ASD=autism spectrum disorder, NT=neurotypical, PCC-iFC=posterior cingulate cortex intrinsic functional connectivity (x=0, y=−53, z=26), VMHC=voxel-mirrored homotopic connectivity, ReHo=regional homogeneity, L=left, R=right. Different sites in ABIDE are color-coded on the top left. Data are shown as residuals obtained after regressing out mean framewise displacement and age effects.

### Additional file 8

**Title:** Stability of sex-by-diagnosis interaction effect

**File format:** tif

**Description:** Inter-site stability of the sex-by-diagnosis interaction pattern was assessed after extracting group means at the mask corresponding to the clusters showing a significant interaction in the discovery analyses and then deriving the group mean when leaving one acquisition site out at the time. The pattern of results was unchanged. Different sites in ABIDE are color-coded in the legend on the right. Due to processing failure of two subjects for VMHC, the sample size comprised 1017 subjects. Abbreviations: ASD=autism spectrum disorder, NT=neurotypical, PCC-iFC=posterior cingulate cortex intrinsic functional connectivity (x=0, y=−53, z=26), VMHC=voxel-mirrored homotopic connectivity, ReHo=regional homogeneity, L=left, R=right. Different sites in ABIDE are color-coded on the top left.

### Additional file 9

**Title:** Robustness to nuisance corrections of main effects of diagnosis and sex

**File format:** tif

**Description:** Cluster-level replication of the results emerging from the voxel-wise discovery analyses in the ABIDE dataset preprocessed using CompCor for the statistically significant main effects of diagnosis (9a) and sex (9b) after preprocessing with GSR and with ICA-AROMA. The second column on the left shows the clusters (*Z*>3.1, *P*<0.01, corrected) with significant diagnostic and sex effects for posterior cingulate cortex intrinsic functional connectivity (PCC-iFC), voxel-mirrored homotopic connectivity (VMHC), and regional homogeneity (ReHo). Results are overlaid on inflated brain maps generated by BrainNet Viewer. The bar plots represent the residual means resulting from regressing out diagnosis or sex effects depending on the desired main effect from the cluster means. 9a) The ABIDE GSR and ABIDE ICA-AROMA columns illustrate, for each of these R-fMRI indices, the diagnostic group mean pattern across clusters with a diagnostic effect size of *η*_*p*_^*2*^≥ 0.01. Color codes: Red=ASD; GreeN=NT. <u>PCC-iFC</u>: bilateral paracingulate cortex and frontal pole (PCG/FP), superior lateral occipital cortex (sLOC), temporal occipital fusiform cortex and occipital fusiform gyrus (TOFC/OFC); <u>VMHC</u>: bilateral posterior cingulate gyrus and precuneus (PCC/Prec), PCG/FP; <u>ReHo</u>: PCG/FP, central operculum and insula (CO/Ins). ABIDE GSR: 7 out of 7 main effects of diagnosis replicated (100%). ABIDE ICA-AROMA: 7 out of 7 main effects of diagnosis replicated (100%) 9b) The ABIDE GSR and ABIDE ICA-AROMA columns illustrate, for each of these R-fMRI indices, the sex group mean pattern across clusters with an effect size of *η*_*p*_^*2*^≥ 0.01. Color codes: Blue=males; Pink=females. <u>PCC-iFC</u>: bilateral sLOC, middle frontal gyrus (MFG), bilateral PCC/Prec, bilateral PCG/FP; <u>VMHC</u>: bilateral PCC/Prec, bilateral anterior cingulate cortex (ACC); <u>ReHo</u>: bilateral PCC, angular gyrus and lateral occipital cortex (AnG/LOC). ABIDE GSR: 10 out of 10 main effects of diagnosis replicated (100%). ABIDE ICA-AROMA: 10 out of 10 main effects of diagnosis replicated (100%). Due to processing failure of two subjects for VMHC, the sample size comprised 1017 subjects instead of 1019. VMHC data are shown as residuals obtained after regressing out mean framewise displacement and age effects. Abbreviations: ASD=autism spectrum disorder, NT=neurotypical, M=males, F=females, CompCor=component base noise reduction, GSR=Global Signal Regression, ICA-AROMA=independent component analysis – automatic removal of motion artifacts, PCC-iFC=posterior cingulate cortex intrinsic functional connectivity (x=0, y=−53, z=26), VMHC=voxel-mirrored homotopic connectivity, ReHo=regional homogeneity, L=left, R=right, R+=replication based on same direction of results and *η*_*p*_^*2*^≥ 0.01, R−=non-replication of results (displayed in gray plots).

### Additional file 10

**Title:** Replicability of main effects of diagnosis and sex

**File format:** tif

**Description:** Cluster-level replication of the results emerging from the voxel-wise analyses in the ABIDE dataset for main effects of diagnosis (10a) and sex (10b) in the Gender Explorations of Neurogenetics and Development to Advance Autism Research (GENDAAR) and the EU-AIMS Longitudinal European Autism Project (LEAP) samples. The ABIDE column on the left shows the clusters (*Z*>3.1, *P*<0.01, corrected) with significant diagnostic and sex effects for posterior cingulate cortex intrinsic functional connectivity (PCC-iFC), voxel-mirrored homotopic connectivity (VMHC), and regional homogeneity (ReHo). No significant effects were detected for degree centrality and fractional amplitude of low frequency fluctuations. Results are overlaid on inflated brain maps generated by BrainNet Viewer. The bar plots represent the residual means resulting from the linear Gaussian regression for each group. 10a) The GENDAAR and EU-AIMS LEAP columns illustrate, for each R-fMRI index, the diagnostic group mean pattern across clusters with a diagnostic effect size of *η*_*p*_^*2*^ ≥ 0.01. Color codes: Red=ASD; Green=NT. <u>PCC-iFC</u>: bilateral paracingulate cortex and frontal pole (PCG/FP), superior lateral occipital cortex (sLOC), temporal occipital fusiform cortex and occipital fusiform gyrus (TOFC/OFC); <u>VMHC</u>: bilateral posterior cingulate gyrus and precuneus (PCC/Prec), PCG/FP; <u>ReHo</u>: PCG/FP, central operculum and insula (CO/Ins). <u>GENDAAR</u>: 6 out of 7 main effects of diagnosis replicated (86%); <u>EU-AIMS LEAP</u>: 2 out of 7 main effects of diagnosis replicated (29%). 10b) The GENDAAR and EU-AIMS LEAP columns illustrate, for each of these R-fMRI index, the sex group mean pattern across clusters with an effect size of *η*_*p*_^*2*^≥ 0.01. Color codes: Blue=males, Pink=females. <u>PCC-iFC</u>: bilateral sLOC, middle frontal gyrus (MFG), bilateral PCC/Prec, bilateral PCG/FP; <u>VMHC</u>: bilateral PCC/Prec, bilateral anterior cingulate cortex (ACC); <u>ReHo</u>: bilateral PCC, angular gyrus and lateral occipital cortex (AnG/LOC). <u>GENDAAR</u>: 5 out of 10 main effects of sex replicated (50%); <u>EU-AIMS LEAP</u>: 8 out of 10 main effects of sex replicated (80%). *Due to processing failure of two subjects for VMHC, the sample size comprised 1017 subjects instead of 1019. Abbreviations: ASD=autism spectrum disorder, NT=neurotypical, M=males, F=females, PCC-iFC=posterior cingulate cortex intrinsic functional connectivity (x=0, y=−53, z=26), VMHC=voxel-mirrored homotopic connectivity, ReHo=regional homogeneity, L=left, R=right, R+=replication based on same direction of results and *η*_*p*_^*2*^≥ 0.01, R−=non-replication of results.

