## Supplementary Material -Additional file 1 for "Towards robust and replicable sex differences in the intrinsic brain function of autism"

***equal contribution**

**Supplemental Methods**

**1.Discovery sample: ABIDE I and II**

**1.1 Inclusion and exclusion criteria**

We selected neurotypical (NT) data from individuals without history of any psychiatric disorders (other than specific phobias in neurotypical children), nor of psychotropic medication use. For the autism datasets, autism diagnosis was determined by clinician’s consensus supported by either one or both ‘gold-standard’ diagnostic instruments, i.e., an Autism Diagnostic Observation Schedule, ADOS [1] and/or the Autism Diagnostic Interview-Revised, ADI-R [2] in all sites but two (UCD and Stanford sites only used diagnostic cut-offs of ADOS and/or ADI-R for inclusion). The corresponding MRI data were included in our discovery analyses based on the following selection criteria and visualized in Additional file 4: **1)** datasets from sites providing both male and female datasets (referred to as an individual’s imaging data) in at least ten subjects per sex/diagnostic group at age 7-18 years. This age range included the most represented ages across sites, i.e., ages that were present in more than three data collections; **2)** datasets from sites reporting full-scale IQ (FIQ) in at least 75% of individuals per sex/diagnostic group — when missing, FIQ data were estimated by either averaging available performance and verbal IQ scores per sex/diagnostic group or imputing FIQ scores by using the mean per sex/diagnostic group at a given site; **3)** datasets that had more than 95% full-brain coverage; **4)** datasets that successfully completed brain image co-registration and transformation to standard space; **5)** datasets with FIQ scores 2.5 standard deviation (SD) within the mean of the group defined by steps 1 to 4 (mean=109, SD=15.7); **6)** datasets with mean framewise displacement (mFD) [3] within three times the interquartile range (IQR) + the third quartile (Q3) of the group defined by steps 1 to 5 (i.e., mFD <0.39mm; this decision was made due to the non-normal distribution of mFD); **7)** *within* and *across* each site, we assessed diagnostic group age-mean matching by excluding the individual dataset with the highest and lowest age year at each group until group mean matching was reached; **8)** following these steps, males and females within diagnostic groups were matched for FIQ and mFD *within* and *across* sites. We opted not to match between diagnostic groups (i.e., autism and NT), as it would have further limited sample size. Additionally, matching diagnostic groups on FIQ may result in non-representative samples across groups and inclusion of potential confounds (Dennis *et al.*, 2009); **9)** at each step, any sites with less than three individual datasets per diagnostic/sex group were excluded. As a result of this stringent selection process, the final ABIDE I and II sample of N=1,019 included N=82 females with autism, N=362 males with autism, N=166 neurotypical females (NT F), and N=409 neurotypical males (NT M).

**1.2. Measures of autism severity**

Given its specificity [1,4], the Autism Diagnostic Observation Schedule (ADOS) was used to assess autism severity. Total calibrated severity scores (CSS; [5]) were available in the ABIDE I and II data repositories. The CSS range from 1 to 10, with higher scores indicating more severe autism symptom severity. CSS were developed by Gotham et al. [5] to allow comparability across ADOS modules 1–3 which vary by age and language abilities. All ABIDE I data were collected using the ADOS-G [1] and total CSS were computed post-hoc using the Gotham *et al.*, (2009) guidelines in N=218 (92%) of the autism datasets selected for this study (N=9 sites; KKI_1, NYU, OHSU, PITT, SDSU, STANFORD,UCLA_1, UM_1, YALE). Total CSS were available in all N=207 autism datasets selected for this study from ABIDE II. Among them, N=60 (N=7 sites; GU, KKI_1, NYU, OHSU, SDSU, UCD, UCLA_1) were obtained from scores collected with ADOS-G and converted to CSS based on Gotham’s guidelines [5]. The remaining N=149 total CSS (N=8 sites; GU, KKI_1, KKI_2, NYU, OHSU, SDSU, UCD, UCLA_1) were obtained based on ADOS-2 administrations. As a note, across the two ADOS editions, Module 3 was the most used (N=200 in ABIDE I and N=185 in ABIDE II), followed by Module 4 (N=19 in ABIDE I and N=22 in ABIDE II) and Module 2 (N=3 in ABIDE I and N=1 in ABIDE II). Total CSS were used to explore brain behavior relationship. Similar analyses were also conducted with the scaled ADOS subscores for social affect (SA) and restricted repetitive behaviors (RRB) scales. CSS for the subscales SA and RRB scales [4] were not available in ABIDE. These analyses were repeated covarying for ADOS modules 3 and 4 categorically while excluding four individuals where ADOS scores had been collected using Module 2 (due to low number). Results were substantially similar across both analyses

**2. Replication samples**

Both independent samples – the EU-AIMS Longitudinal European Autism Project (EU-AIMS LEAP) and the Gender Explorations of Neurogenetics and Development to Advance Autism Research (GENDAAR) - were based on multisite datasets collected with harmonized MRI and behavioral assessment protocols across sites within each data collection, separately (5 sites for EU-AIMS LEAP and 4 sites for GENDAAR).

As described elsewhere [6], inclusion criteria for the autism group in the **EU-AIMS LEAP** were an existing clinical autism spectrum disorder (ASD) or equivalent diagnosis according to DSM-IV/ICD-10 or DSM-5 criteria. Autism Diagnostic Interview-Revised (ADI-R) [2] and the Autism Diagnostic Observation Schedule, 2^nd^ edition (ADOS-2) also characterizes the sample. Individuals with a clinical ASD diagnosis who did not reach diagnostic cut-offs on these instruments were not excluded. Individuals with psychosis or bipolar disorder were excluded. For the purpose of this study only datasets from individuals labeled as NT and without history of depression, anxiety, and of psychoactive medication use were included.

For **GENDAAR**, and as described elsewhere [7–9], participants in the autism group were required to have a prior clinical diagnosis of ASD that was confirmed by research-reliable clinicians using the ADOS-2 [10] and/or the ADI-R [2] – notably a protocol was in place to maintain research reliability over time across evaluators between and within sites [8]. Absence of known neurodevelopmental diagnosis and having a first-or second-degree relative with ASD, and no evidence of elevated ASD traits (total t scores < 65 on the parent-report version of the Social Responsiveness Scale, Second Edition [SRS-2]; [11]), were required for NT controls. Further, unaffected siblings of individuals with autism were also excluded. Exclusion criteria for both groups were prematurity, a genetic, neurological or psychiatric comorbidity. These encompass Fragile X syndrome, epilepsy, brain injury, pre-/peri-natal birth injury, nutritional or psychological deprivation, visual or auditory impairment after correction, sensorimotor difficulties, use of any benzodiazepine, barbiturate or anti-epileptic medication, pregnancy and tic disorders.

For both these replication samples, only datasets with more than 95% full-brain coverage after successful anatomical and functional registration were included. Only datasets with age, FIQ in the same range as that of the discovery ABIDE sample (i.e., age=7-18 years; FIQ=70-148) and, similarly, mFD<0.39 were included. For the GENDAAR sample only one member of a sibling pair was included. For a comparison of demographic and clinical information between ABIDE, GENDAAR and EU-AIMS LEAP, see Additional file 3: Tables S3, S4 and S5. Briefly, there were significant group mean differences in diagnostic group means for ADOS total CSS (EU-AIMS LEAP < ABIDE = GENDAAR), age (ABIDE < EU-AIMS LEAP = GENDAAR) between the autism groups. In regard to FIQ, significant differences were noted between independent samples, albeit only for the NT groups (ABIDE > EU-AIMS LEAP = GENDAAR).

**3. R-fMRI measures**

*Seed-based Correlation Analysis* (SCA) was carried out by extracting the mean time series from a spherical region-of-interest mask (8mm in diameter) centered in PCC using the seed location: x=0, y=-53, z=26 previously defined by Andrews-Hanna et al., (2007) [12] and used in Di Martino et al., (2014) [13] and Floris et al., (2018) [14]. Pearson’s correlation coefficient was calculated between the PCC time series and each voxel in the brain before being Fisher’s z-transformed. PCC-iFC is commonly investigated in autism [13–18] and there are known sex differences in the default network in neurotypicals [8,19–22].

*Voxel-Mirrored Homotopic Connectivity (VMHC)* [23] is the Pearson’s correlation between each voxel and its geometrically corresponding symmetric counterpart in the opposite hemisphere. Spatial transformation parameters were based on a registration to a symmetric MNI template [24] to increase spatial correspondence between homotopic voxels. Correlation coefficients were standardized by applying a Fisher’s r-to-z-transformation. Atypical cross-hemispheric homotopic connectivity has previously been reported in autism [13,14,25–28] and shown to differ across the sexes in neurotypicals [23,25].

*Regional Homogeneity (ReHo)* [29] is a measure of regional coherence between neighboring fMRI time series. It is based on the Kendall’s coefficient of concordance [30] between a voxel’s time series and its 26 adjacent neighbors. Subject-level maps were transformed into subject-level z-score maps. Previous studies have reported differences in local connectivity in autism [13,14,31–36] and between the sexes in NT [37–39].

*Network Degree Centrality (DC)* [40] is a measure of local network connectivity. To be consistent with prior studies [13,14,41], here, it is based on a given voxel’s sum of significant connections with corresponding *p* < 0.001. DC was calculated based on a study-specific functional volume mask based on voxels (in MNI space) present in at least 90% of subjects and further constrained by a 25% gray matter (GM) probability mask. Voxel-size was down-sampled to 4mm^3^ to reduce computational intensity. Voxel-based graphs were then generated by computing the Pearson’s correlation of each voxel’s extracted time series with every other voxel’s extracted time series within the study-specific mask. A significance threshold of *p* < 0.001 was applied resulting in a binary, undirected adjacency matrix. DC was then computed by counting the number of significant connections in the adjacency matrix. Subject-level DC-maps were standardized using z-score transformations. Degree centrality is among the graph theoretical measures most commonly used in R-fMRI studies in autism [13,14,42–45] and has been shown to differ across the sexes in NT [40,41].

*Fractional Amplitude of Low Frequency Fluctuations (fALFF)* [46] is a frequency domain metric representing the relative contribution of specific oscillations to the entire frequency range. It is based on the ratio of the amplitudes of fluctuations in the 0.01-0.1 Hz frequency range to the sum of amplitudes in the entire frequency spectrum. No temporal filtering was applied, because the data were analyzed in the frequency domain. Fractional ALFF maps at the subject-level were transformed into subject-level z-score maps. Previous studies have shown differences in fALFF in individuals with autism compared to neurotypical controls [13,14,45,47–49] and between males and females in NT [37,41,46].

**4. Preprocessing Pipeline**

**4.1 Structural preprocessing**

1) Skull-stripping: T1-weighted images were skull-stripped using FSL’s BET [50] command.
2) Tissue segmentation: FSL’s FAST [51] command was used to segment images into GM, white matter (WM) and cerebrospinal fluid (CSF). Probability thresholds were 0.96 for WM and CSF, and 0.7 for GM.
3) Spatial normalization: images (with skull-on) were normalized to MNI152 stereotactic space (2mm^3^ isotropic) with linear and non-linear registrations using ANTs [52]. For the calculation of VMHC, spatial normalization was done by registering to a symmetrical template.

**4.2 Functional preprocessing**

1) Slice time correction: the AFNI command *3dTshift* was used to correct for differences in acquisition time between the slices using the specific parameters for each site based on their acquisition protocols.
2) Motion realignment: motion correction was performed in two steps. First using the AFNI command *3dvolreg* by each functional volume was co-registered to the (un-aligned) mean functional image. In a second step, a new functional mean image based on the aligned images was used as the reference image. At this second stage, motion parameters based on the Friston 24-Parameter Model (six motion parameters, their values of preceding volumes, 12 squared values of these items) were calculated along with mean framewise displacement (mFD) [3].
3) Skull-stripping: skull was removed using the AFNI command *3dAutomask*.
4) Mean-based intensity normalization: all images were scaled with a factor of 10.000.
5) Nuisance signal regression differed across the discovery and robustness analyses. Across all pipelines 24 motion parameters based on Friston 24-Parameter Model were regressed out along with linear and quadratic trends. For the discovery analysis, the component-based noise correction (CompCor) was used [53]. For robustness analyses two different regression methods were used: Independent Component Analysis - Automatic Removal of Motion Artifacts (ICA-AROMA) [54] and global signal regression (GSR). Details of each of these methods are addressed below.
6) Temporal filtering: band-pass filtering (0.01-0.1 Hz) was done using the AFNI command *3dBandpass* for all R-fMRI derivatives other than fALFF.
7) Registration: functional-to-anatomical co-registration was achieved by Boundary Based Registration (BBR) [55] using FSL FLIRT. Spatial normalization of functional EPIs to MNI152 space was done by applying linear and non-linear transforms from ANTs.
8) ReHo, fALFF, and SCA of PCC were calculated in native space, before being transformed into MNI152 space. DC was calculated in MNI152 space. As above, VMHC was calculated based on smoothed data in symmetric MNI152 space.
9) Spatial filtering: Derivatives (fALFF, ReHo, SCA, DC) were smoothed with a 3D Gaussian kernel (FWHM=6mm) after computing and registering each derivative. VMHC was spatially filtered (FWHM=6mm) prior to its calculation and registration.

**5. ComBat**

Given that the discovery and replication samples were all multisite, to account for site and collection time variability we applied ComBat <https://github.com/brentp/combat.py>) (23). ComBat harmonization is a statistical technique originally designed for genomic studies to correct unwanted non-biological variability across multiple batches of gene expression microarray experiments [56], also called “batch effects.” In genomic studies, these batch effects are systematically introduced when new samples are added to an existing dataset of arrays or in multiple studies that make use of microarrays data across different labs. In the context of MRI images, the batch effects can be introduced by differences in the scanner manufacturers, differences in data collection timing and sites. Recent studies demonstrated that ComBat is a robust method to correct confounding differences on multi-site DTI [57], cortical thickness [58] and fMRI [59] datasets. ComBat uses a parametrical and non-parametrical empirical Bayes framework defined by the following location and scale (L/S) batch adjustment model:

$$Y_{ijv}=a_{v}+X\beta_{v}+\gamma_{iv}+\delta_{iv}\varepsilon_{ijv}$$

where $Y_{ijv}$ represents the expression value for voxel $v$ for sample $j$ from site $i$, $a_{v}$ is the Z-score value for voxel $v$, $X$ represents the design matrix for nuisance signals, and $\beta_{v}$ is the vector of regression coefficients corresponding to $X$. The error term $\varepsilon_{ijv}$ is assumed to have a normal distribution with zero mean and variance $\sigma_{v}^{2}$. The parameters $\gamma_{iv}$ and $\delta_{iv}$ are the additive and multiplicative site effects $i$ for voxel $v$, respectively, and both are estimated empirically by assuming that all voxels share the same common distribution [56,57].

**6. Robustness analyses**

In discovery analyses on the ABIDE datasets*,* nuisance signal regression was performed using component-based noise correction (CompCor) [53] to remove physiological noise. Using CompCor the first 5 principal components from a combined WM/CSF mask are regressed out at each individual general linear model. To assess the robustness of the results obtained in discovery analyses to changes in preprocessing approaches, we repeated the analyses replacing the CompCor step – with either ICA-AROMA [54] or GSR [60,61] as nuisance regressions approaches. All other structural and functional preprocessing steps remained unchanged. For both the ICA-AROMA and GSR regression pipelines, WM and CSF signals were also regressed.

**ICA-AROMA**. It aims to reduce motion-induced signals variation in R-fMRI. This technique largely preserves the autocorrelation structure of the fMRI time-series and it also avoids the reduction of the temporal degrees of freedom, which are known drawback of scrubbing and regressing of motion volumes techniques [54]. ICA-AROMA steps include: 1) automatic estimation and extraction of independent components (ICs) implemented in the FSL’s probabilistic ICA tool, MELODIC [62], 2) automatic motion-related ICs classification based on a combination of four discriminative features that represent motion artifacts: high-frequency content features, realignment parameters and spatial features consisting of edge and CSF fraction metrics, and 3) removal of the classified ICs from the fMRI data using linear regression (Pruim et al., 2015).

**GSR**. Global signal regression (GSR) involves the averaging of all voxel time-series within a gray matter / whole brain mask and its inclusion in the general linear model analysis as a nuisance regressor. For this study, GSR was computed based on the gray matter mask available in CPAC. As a time-varying spatial average, GSR is thought to improve the detection of neuronal signals by removing other sources of global variance (e.g., motion and respiratory-related artifacts) [63,64]. However, GSR has also been reported to alter functional connectivity maps by shifting the distribution of iFC values and introducing negative correlations [65]. As the discussion on whether to use the global signal as a nuisance regressor has been ongoing in the fMRI community, we opted to also include it as an approach for the robustness strategy.

All preprocessing pipelines were conducted using the Configurable Pipeline for the Analysis of Connectomes (C-PAC, <http://fcp-indi.github.com/C-PAC/>). CPAC version 0.3.9 was used for all analyses with the exception of those focusing on ICA-AROMA, which was not implemented until V1.30. Regression-testing performed at the time of each CPAC version release was used to confirm that there were no appreciable changes between CPAC versions in any key intermediates (e.g., nuisance covariates) or derived data calculated (i.e., concordance correlation coefficient > 0.98 against an established reference output benchmark).
