## Supplementary Table - Additional file 3 for "Towards robust and replicable sex differences in the intrinsic brain function of autism"

**Table S1. Characterization of EU-AIMS LEAP sample**

| EU-AIMS LEAP | Sites^a^ | ASD M (N=133) | ASD F (N=43) | NT M (N=85) | NT F (N=48) |  |  |
| --- | --- | --- | --- | --- | --- | --- | --- |
|  | **N** | **Mean (SD)**  **[Range]** | **Mean (SD)**  **[Range]** | **Mean (SD)**  **[Range]** | **Mean (SD)**  **[Range]** | **Statistics** | **Post-hoc** |
| Age | 5 | 14.2 (3.0)  [7.5- 18.9] | 13.3 (3.4)  [7.1-18.9] | 13.9 (3.0)  [7.6- 18.8] | 13.9 (3.8)  [6.9- 18.6] | *F_(3)_*=0.83, *p*=0.48 |  |
| Full-Scale IQ^b^ | 5 | 103 (16)  [72-148] | 101 (15.9)  [70.4-131] | 108 (15)  [71.7-140] | 107 (13.2)  [72.7-133] | *F*_(3)_=2.59, *p*=0.05 | (ASD M=ASD F) < (NT M=NT F) |
| Verbal IQ^b^ | 5 | 102 (16.9)  [67-144] | 100 (15)  [67-136] | 106 (15.4)  [73-142] | 105 (14.9)  [65-140] | *F*_(3)_=2.11, *p*=0.1 |  |
| Performance IQ^b^ | 5 | 105 (17.9)  [59-150] | 101 (18.7)  [58-133] | 109 (17.7)  [61-139] | 107 (14.4)  [70-139] | *F*_(3)_=1.85, *p*=0.14 |  |
| Mean FD | 5 | 0.13 (0.08)  [0.02-0.35] | 0.1 (0.07)  [0.04-0.33] | 0.1 (0.07)  [0.03-0.31] | 0.09 (0.07)  [0.02-0.36] | *H_(3)_*=15.88, *p*=0.001 | (ASD M>ASD F) < (NT M=NT F) |
| ADI-R |  |  |  |  |  |  |  |
| Social | 5 | 17.2 (6.3)  [1-29] | 16.1 (7.8)  [1-27] | - | - | *t_(62)_*=0.84, *p*=0.41 |  |
| Communication | 5 | 13.7 (5.5)  [2-26] | 12.8 (5.6)  [1-24] | - | - | *t_(71)_*=0.93, *p*=0.36 |  |
| RRB | 5 | 4.5 (2.8)  [0-12] | 4.4 (2.7)  [0-10] | - | - | *t_(74)_*=0.23, *p*=0.82 |  |
| ADOS-2 |  |  |  |  |  |  |  |
| Social-Affect CSS^c^ | 4 | 6.2 (2.6)  [1-10] | 5.4 (2.6)  [1-10] | - | - | *t_(70)_*=1.65, *p*=0.1 |  |
| RRB CSS^c^ | 4 | 4.6 (2.7)  [1-10] | 4.6 (2.5)  [1-9] | - | - | *t_(77)_*=-0.01, *p*=0.99 |  |
| CSS total^d^ | 4 | 5.5 (2.8)  [1-10] | 4.5 (2.6)  [1-9] | - | - | *t_(76)_*=2.08, *p*=0.04 |  |
|  |  | **N** | **N** |  |  | **Statistics** | **Post-hoc** |
| Comorbidity | 4 | 52^e^ | 29^f^ | - | - | *χ^2^_(1)_*=9.4, *p*=0.002 |  |
| Psychoactive Meds | 4 | 102 | 25 | - | - | *χ^2^_(1)_*=4.7, *p*=0.03 |  |

Abbreviations: *ADI-R* = Autism Diagnostic Interview-Revised; *ADOS-2* = Autism Diagnostic Observation Schedule-2; *ASD* = Autism Spectrum Disorder; *CSS* = Calibrated Severity Score*; F* = females; *IQ* = intellectual quotient; *M* = males; *Mean FD* = mean framewise displacement (Jenkinson *et al.*, 2002); *NT* = neurotypical; RRB= restricted repetitive behaviors. ^a^EU-AIMS LEAP data collections: Kings College London, UK, Cambridge University, UK; Donders Institute Nijmegen, Netherlands; University of Utrecht, Netherlands; ZI Mannheim, Germany;  ^b^FIQ, VIQ and PIQ were assessed using the WASI or WISC / WAIS;  ^c^ Social-Affect & RRB Calibrated Severity Scores computed based on (Hus *et al.*, 2014) for Module 3 and (Hus and Lord, 2014) for Module 4; ^d^Total Calibrated Severity Score computed based on (Gotham *et al.*, 2009) for Module 3 and based on (Hus and Lord, 2014) for Module 4; ^e^ADHD (N=49); anxiety disorder (N=5); depression (N=5). ^f^ADHD (N=21); anxiety disorder (N=3); depression (N=2). The three group means were compared with ANOVA tests (or Kruskal-Wallis test in the case of non-parametric mean FD) followed by post-hoc pairwise t-test comparisons (or Mann-Whitney U-tests in the case of non-parametric mean FD) when statistically significant (significance cut-off set at p<0.05).

**Table S2. Characterization of GENDAAR sample**

| GENDAAR | Sites^a^ | ASD M (N=43) | ASD F (N=44) | NT M (N=56) | NT F (N=53) |  | | |
| --- | --- | --- | --- | --- | --- | --- | --- | --- |
|  | **N** | **Mean (SD)**  **[Range]** | **Mean (SD)**  **[Range]** | **Mean (SD)**  **[Range]** | **Mean (SD)**  **[Range]** | **Statistics** | **Post-hoc** | |
| Age | 4 | 13.4 (3.0)  [8.2- 17.9] | 13.6 (2.7)  [8.2-18.0] | 13.7 (2.7)  [8.4- 17.8] | 13.7 (2.8)  [8.2- 17.9] | *F_(3)_*=1.22, *p*=0.93 |  | |
| Full-Scale IQ^b^ | 4 | 101 (17.2)  [71-139] | 102 (19.9)  [70-145] | 112 (14.8)  [79-143] | 111 (14.0)  [83-139] | *F*_(3)_=5.86, *p*<0.001 | (ASD M=ASD F) < (NT M=NT F) | |
| Verbal IQ^c^ | 4 | 102 (20.5)  [52-152] | 103 (19.2)  [63-147] | 112 (17.3)  [74-159] | 110 (14.0)  [84-138] | *F*_(3)_=3.8, *p*=0.011 |  | |
| Performance IQ^d^ | 4 | 101 (15.2)  [61-136] | 101 (19.6)  [66-143] | 111 (13.2)  [85-136] | 109 (13.7)  [84-139] | *F*_(3)_=2.57, *p*=0.06 |  | |
| Mean FD | 4 | 0.16 (0.1)  [0.03-0.39] | 0.16 (0.11)  [0.03-0.37] | 0.13 (0.09)  [0.03-0.36] | 0.11 (0.1)  [0.02-0.39] | *H_(3)_*=11.6, *p*=0.008 | (ASD M=ASD F) < (NT M=NT F) | |
| ADI-R |  |  |  |  |  |  | | |
| Social | 4 | 19.6 (5.6)  [5-27] | 19.0 (6.0)  [1-30] | - | - | *t_(82)_*=0.49, *p*=0.63 | |  |
| Communication | 4 | 17.0 (4.2)  [8-26] | 15.5 (4.6)  4-24] | - | - | *t_(81)_*=1.56, *p*=0.12 | |  |
| RRB | 4 | 6.0 (2.6)  [1-12] | 5.9 (3.1)  [0-12] | - | - | *t_(80)_*=0.15, *p*=0.88 | |  |
| ADOS-2 |  |  |  |  |  |  | | |
| Social-Affect | 4 | 10.1 (4.8)  [0-19] | 7.9 (3.5)  [1-18] | - | - | *t_(59)_*=2.19, *p*=0.03 | | ASD M>ASD F |
| RRB | 4 | 2.9 (1.7)  [0-6] | 2.0 (1.8)  [0-6] | - | - | *t_(71)_*=1.97, *p*=0.05 | | ASD M>ASD F |
| CSS total | 4 | 7.3 (2.4)  [1-10] | 6 (2.3)  [1-10] | - | - | *t_(64)_*=2.37, *p*=0.02 | | ASD M>ASD F |

*ADI-R* = Autism Diagnostic Interview-Revised; *ADOS-2* = Autism Diagnostic Observation Schedule-2; *ASD* = Autism Spectrum Disorder; *CSS* = Calibrated Severity Score*; F* = females; *GENDAAR*= Gender Explorations of Neurogenetics and Development to Advance Autism Research; *IQ* = intellectual quotient; *M* = males; *Mean FD* = mean framewise displacement (Jenkinson *et al.*, 2002); *NT* = neurotypical; RRB= restricted repetitive behaviors. ^a^GENDAAR data collections: The Nelson Laboratory of Cognitive Neuroscience, Boston Children’s Hospital, Harvard Medical School, Boston, MA; the Center on Human Development & Disability, Seattle Children’s Hospital, University of Washington School of Medicine, Seattle, WA; Staglin IMHRO Center for Cognitive Neuroscience, David Geffen School of Medicine, University of California, Los Angeles, CA; ^b^FIQ, VIQ and PIQ (Non-Verbal Reasoning) were assessed using the Differential Ability Scales (DAS II). The three group means were compared with ANOVA tests (or Kruskal-Wallis test in the case of non-parametric mean FD) followed by post-hoc pairwise t-test comparisons (or Mann-Whitney U-tests in the case of non-parametric mean FD) when statistically significant (significance cut-off set at p<0.05).

**Table S3. Comparison between ABIDE vs. EU-AIMS LEAP sample**

|  | **ASD M_ABIDE_ vs. ASD M_EU-AIMS LEAP_** |  | **ASD F_ABIDE_ vs. ASD F _EU-AIMS LEAP_** |  | **NT M_ABIDE_ vs. NT M _EU-AIMS LEAP_** |  | **NT F_ABIDE_ vs. NT F _EU-AIMS LEAP_** |  |
| --- | --- | --- | --- | --- | --- | --- | --- | --- |
| **Age** | *t*_(208)_=-7.9, *p*<0.001 | **A < E** | *t*_(71)_=-2.6, *p*=0.01 | **A < E** | *t*_(112)_=-6.2, *p*<0.001 | **A < E** | *t*_(57)_=-4.5, *p*<0.001 | **A < E** |
| **FIQ** | *t*_(238)_=1.8, *p*=0.08 | A = E | *t*_(87)_=0.9, *p*=0.34 | A = E | *t*_(109)_=2.5, *p*=0.01* | **A > E** | *t*_(74)_=3.3, *p*=0.001* | **A > E** |
| **VIQ** | *t*_(254)_=2.9, *p*<0.01* | **A > E** | *t*_(99)_=1.6, *p*=0.12 | A = E | *t*_(115)_=4.4, *p*<0.001* | **A > E** | *t*_(80)_=3.5, *p*<0.001* | **A > E** |
| **PIQ** | *t*_(237)_=0.6, *p*=0.55 | A = E | *t*_(84)_=0.8, *p*=0.44 | A = E | *t*_(110)_=-0.11, *p*=0.91 | A = E | *t*_(76)_=1.1, *p*=0.29 | A = E |
| **Mean FD** | *U*=21380, *p*=0.06 | A = E | *U*=2189, *p*=0.03* | **A > E** | *U*=16233, *p*=0.34 | A = E | *U*=4136, *p*=0.69 | A = E |
| **ADOS CSS total** | *t*_(183)_=4.8, *p*<0.001* | **A > E** | *t*_(65)_=5.3, *p*<0.001* | **A > E** | - | *-* | - | *-* |
| **ADI-R social** | *t*_(201)_=3.9, *p*<0.001* | **A > E** | *t*_(70)_=2.5, *p*=0.01* | **A > E** | - | *-* | - | *-* |
| **ADI-R comm** | *t*_(202)_=3.4, *p*<0.001* | **A > E** | *t*_(83)_=2.2, *p*<0.03* | **A > E** | - | - | - | - |
| **ADI-R RRB** | *t*_(209)_=5.2, *p*<0.001* | **A > E** | *t*_(84)_=2.7, *p*=0.01* | **A > E** | - | - | - | - |
| *Abbreviations: A =* Autism Brain Imaging Data Exchange (ABIDE)*; ADI-R* = Autism Diagnostic Interview-Revised; *ADI-R comm* = ADI-R total communication subscore; ADI-R RRB = ADI-R total restricted repetitive behaviors subscore; *ADOS* = Autism Diagnostic Observation Schedule; *ASD* = Autism Spectrum Disorder; *CSS* = Calibrated Severity Score*; E=* EU-AIMS Longitudinal European Autism Project (EU-AIMS LEAP); *F* = females; *IQ* = intellectual quotient; *FIQ*= full IQ; *VIQ* = verbal IQ; *PIQ* = performance IQ; *M* = males; *Mean FD* = mean framewise displacement (Jenkinson *et al.*, 2002); *NT* = neurotypical. See Table 1 and Supplemental Table 1 for group means and SD and Supplemental Material for inclusion, exclusion and study selection criteria for these datasets. * indicate statistically significant with two-tailed tests setting a significance cut-off at p<0.05. Due to non-normal distribution of Mean FD, a non-parametric Man-Whitney U test was used in this case. | | | | | | | | |

**Table S4. Comparison between ABIDE vs. GENDAAR sample**

|  | **ASD M_ABIDE_ vs. ASD M_GENDAAR_** |  | **ASD F_ABIDE_ vs. ASD F_GENDAAR_** |  | **NT M_ABIDE_ vs. NT M_GENDAAR_** |  | **NT F_ABIDE_ vs. NT F_GENDAAR_** |  |
| --- | --- | --- | --- | --- | --- | --- | --- | --- |
| **Age** | *t*_(50)_=-3.3, *p*<0.01* | **A < G** | *t*_(88)_=-3.7, *p*<0.001* | **A < G** | *t*_(70)_=-5.1, *p*<0.001* | **A < G** | *t*_(74)_=-5.4, *p*<0.001* | **A < G** |
| **FIQ** | *t*_(52)_=1.8, *p*<0.07 | A = G | *t*_(75)_=0.6, *p*=0.55 | A = G | *t*_(67)_=0.2, *p*=0.87 | A = G | *t*_(77)_=1.3, *p*=0.2 | A = G |
| **VIQ** | *t*_(51)_=1.4, *p*=0.15 | A = G | *t*_(84)_=0.6, *p*=0.54 | A = G | *t*_(66)_=1.0, *p*=0.34 | A = G | *t*_(92)_=1.7, *p*=0.1 | A = G |
| **PIQ** | *t*_(53)_=1.2, *p*=0.23 | A = G | *t*_(85)_=0.6, *p*=0.54 | A = G | *t*_(74)_=-0.5, *p*=0.64 | A = G | *t*_(77)_=0.4, *p*=0.65 | A = G |
| **Mean FD** | *U*=5274, *p*<0.001* | **A < G** | *U*=1649, *p*=0.42 | A = G | *U*=9028, *p*=0.01* | **A < G** | *U*=4067, *p*=0.41 | A = G |
| **ADOS CSS total** | *t*_(35)_=-1.1, *p*=0.23 | A = G | *t*_(55)_=1.9, *p*=0.06 | A = G | - | *-* | - | *-* |
| **ADI-R soc** | *t*_(51)_=0.06, *p*=0.94 | A = G | *t*_(83)_=0.5, *p*=0.62 | A = G | - | *-* | - | *-* |
| **ADI-R comm** | *t*_(54)_=-2.03, *p*=0.05 | A = G | *t*_(93)_=-0.35, *p*=0.73 | A = G | - | - | - | - |
| **ADI-R RRB** | *t*_(51)_=0.01, *p*=0.99 | A = G | *t*_(74)_=-0.13, *p*=0.9 | A = G | - | - | - | - |
| *Abbreviations: A =* Autism brain imaging data exchange (ABIDE)*; ADI-R* = Autism Diagnostic Interview-Revised; *ADI-R comm* = ADI-R total communication subscore; ADI-R RRB = ADI-R total restricted repetitive behaviors subscore; *ADOS* = Autism Diagnostic Observation Schedule; *ASD* = Autism Spectrum Disorder; *CSS* = Calibrated Severity Score*; F* = females; *IQ* = intellectual quotient; *FIQ*= full IQ; *G=* Gender Explorations of Neurogenetics and Development to Advance Autism Research (GENDAAR); *VIQ* = verbal IQ; *PIQ* = performance IQ; *M* = males; *Mean FD* = mean framewise displacement (Jenkinson *et al.*, 2002); *NT* = neurotypical. See Table 1 and Supplemental Table 2 for group means and SD and Supplementary Material for inclusion, exclusion and study selection criteria for these datasets. * indicate statistically significant with two-tailed tests setting a significance cut-off at p<0.05. Due to non-normal distribution of Mean FD, a non-parametric Man-Whitney U test was used in this case. | | | | | | | | |

**Table S5. Comparison between EU-AIMS LEAP vs. GENDAAR sample**

|  | **ASD M_EU-AIMS LEAP_ vs. ASD M_GENDAAR_** |  | **ASD F_EU-AIMS LEAP_ vs. ASD F_GENDAAR_** |  | **NT M_EU-AIMS LEAP_ vs. NT M_GENDAAR_** |  | **NT F_EU-AIMS LEAP_ vs. NT F_GENDAAR_** |  |
| --- | --- | --- | --- | --- | --- | --- | --- | --- |
| **Age** | *t*_(72)_=1.5, *p*=0.14 | E = G | *t*_(80)_=-0.5, *p*=0.62 | E = G | *t*_(127)_=0.4, *p*=0.67 | E = G | *t*_(87)_=0.3, *p*=0.11 | E = G |
| **FIQ** | *t*_(68)_=0.7, *p*=0.48 | E = G | *t*_(82)_=-0.2, *p*=0.83 | E = G | *t*_(120)_=-1.6, *p*=0.11 | E = G | *t*_(96)_=-1.7, *p*=0.1 | E = G |
| **VIQ** | *t*_(62)_=-0.1, *p*=0.88 | E = G | *t*_(81)_=-0.7, *p*=0.48 | E = G | *t*_(109)_=-2.0, *p*=0.04* | **E < G** | *t*_(96)_=1.7, *p*=0.1 | E = G |
| **PIQ** | *t*_(71)_=0.8, *p*=0.44 | E = G | *t*_(85)_=-0.1, *p*=0.89 | E = G | *t*_(133)_=-0.3, *p*=0.79 | E = G | *t*_(97)_=-0.5, *p*=0.64 | E = G |
| **Mean FD** | *U*=2265, *p*=0.04* | **E < G** | *U*=660, *p*=0.02* | **E < G** | *U*=2065, *p*=0.18 | E = G | *U*=1126, *p*=0.32 | E = G |
| **ADOS CSS total** | *t*_(53)_=-3.7, *p*<0.001* | **E < G** | *t*_(76)_=-2.8, *p*<0.01* | **E < G** | - | *-* | - | *-* |
| **ADI-R soc** | *t*_(78)_=-2.3, *p*=0.02* | **E < G** | *t*_(79)_=-2.0, *p*=0.05* | **E < G** | - | *-* | - | *-* |
| **ADI-R comm** | *t*_(90)_=-4.0, *p*<0.001* | **E < G** | *t*_(81)_=-2.5, *p*=0.01* | **E < G** | - | - | - | - |
| **ADI-R RRB** | *t*_(74)_=3.1, *p*<0.01* | **E < G** | *t*_(83)_=2.5, *p*=0.02* | **E < G** | - | - | - | - |
| *Abbreviations: ADI-R* = Autism Diagnostic Interview-Revised; *ADI-R comm* = ADI-R total communication subscore; ADI-R RRB = ADI-R total restricted repetitive behaviors subscore; *ADOS* = Autism Diagnostic Observation Schedule; *ASD* = Autism Spectrum Disorder; *CSS* = Calibrated Severity Score*; E =* EU-AIMS Longitudinal European Autism Project (EU-AIMS LEAP); *F* = females; *IQ* = intellectual quotient; *FIQ*= full IQ;  *G=* Gender Explorations of Neurogenetics and Development to Advance Autism Research (GENDAAR); *VIQ* = verbal IQ; *PIQ* = performance IQ; *M* = males; *Mean FD* = mean framewise displacement (Jenkinson *et al.*, 2002); *NT* = neurotypical. See Supplemental Tables 1 and 2 for group means and SD and supplementary Material for inclusion, exclusion and study selection criteria for these datasets. * indicate statistically significant with two-tailed tests setting a significance cut-off at p<0.05. Due to non-normal distribution of Mean FD, a non-parametric Man-Whitney U test was used in this case. | | | | | | | | |
