## Supplementary figures and images for "Towards robust and replicable sex differences in the intrinsic brain function of autism"

### Additional file 2

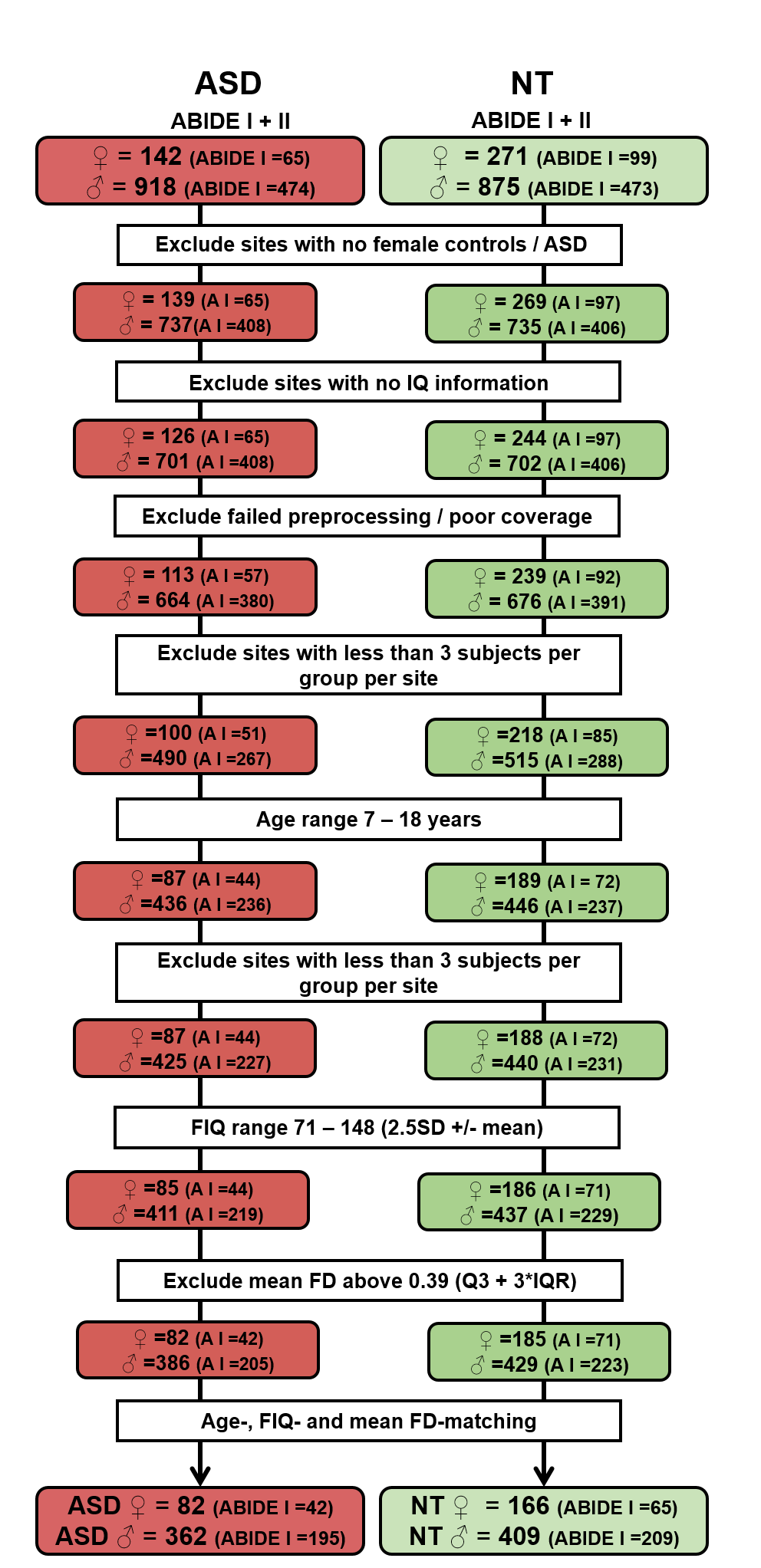

### Additional file 4

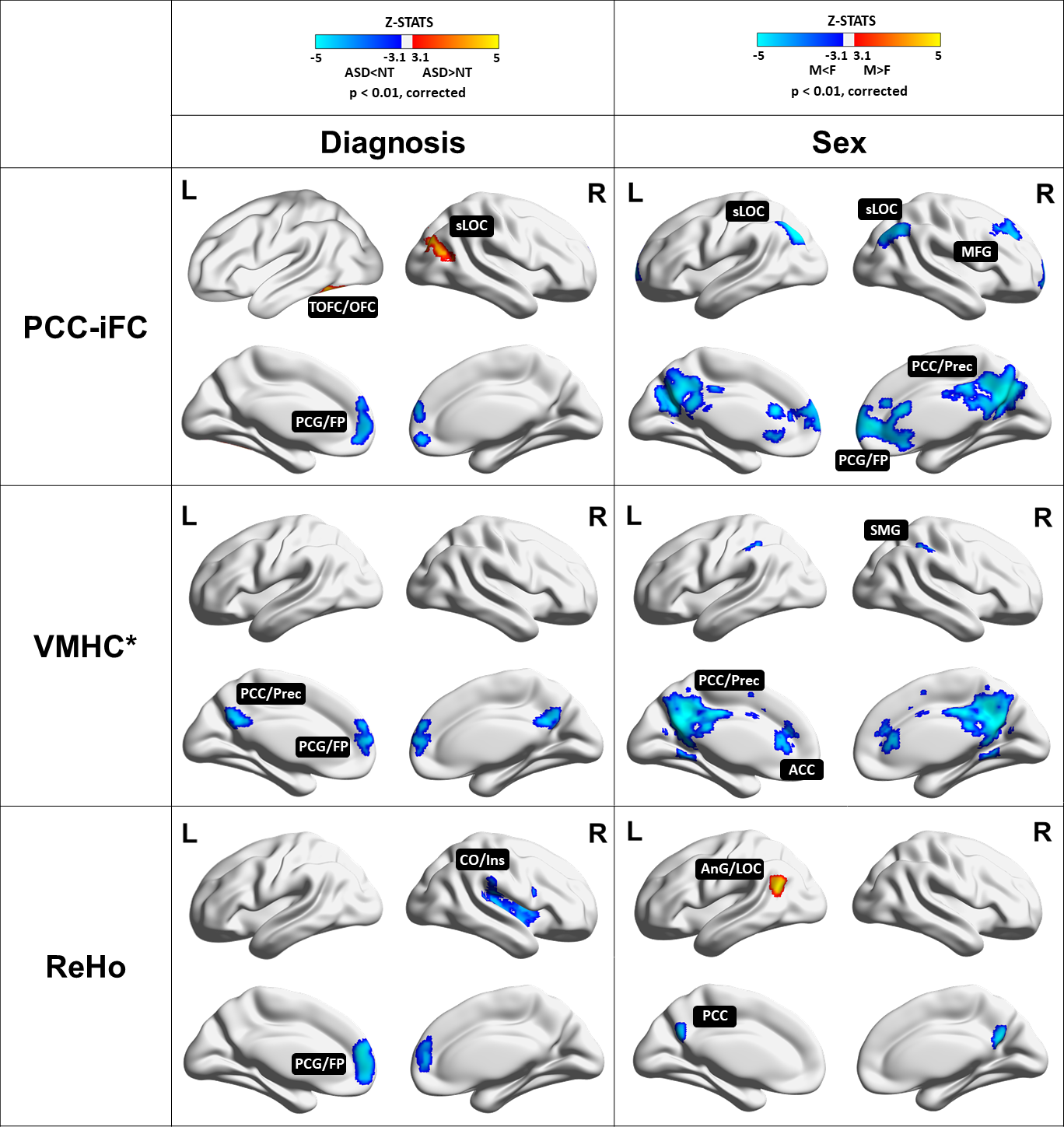

### Additional file 6

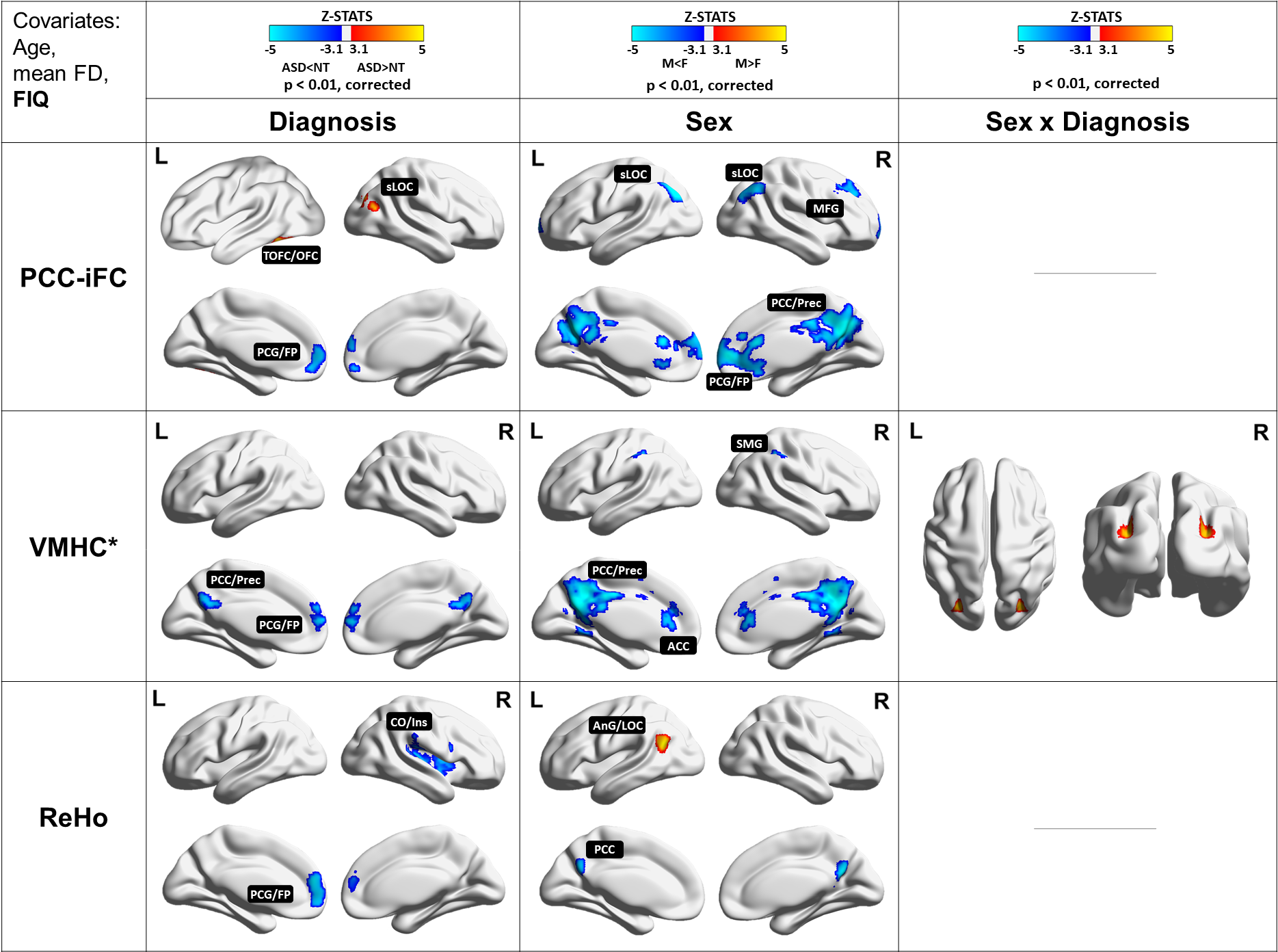

### Additional file 7

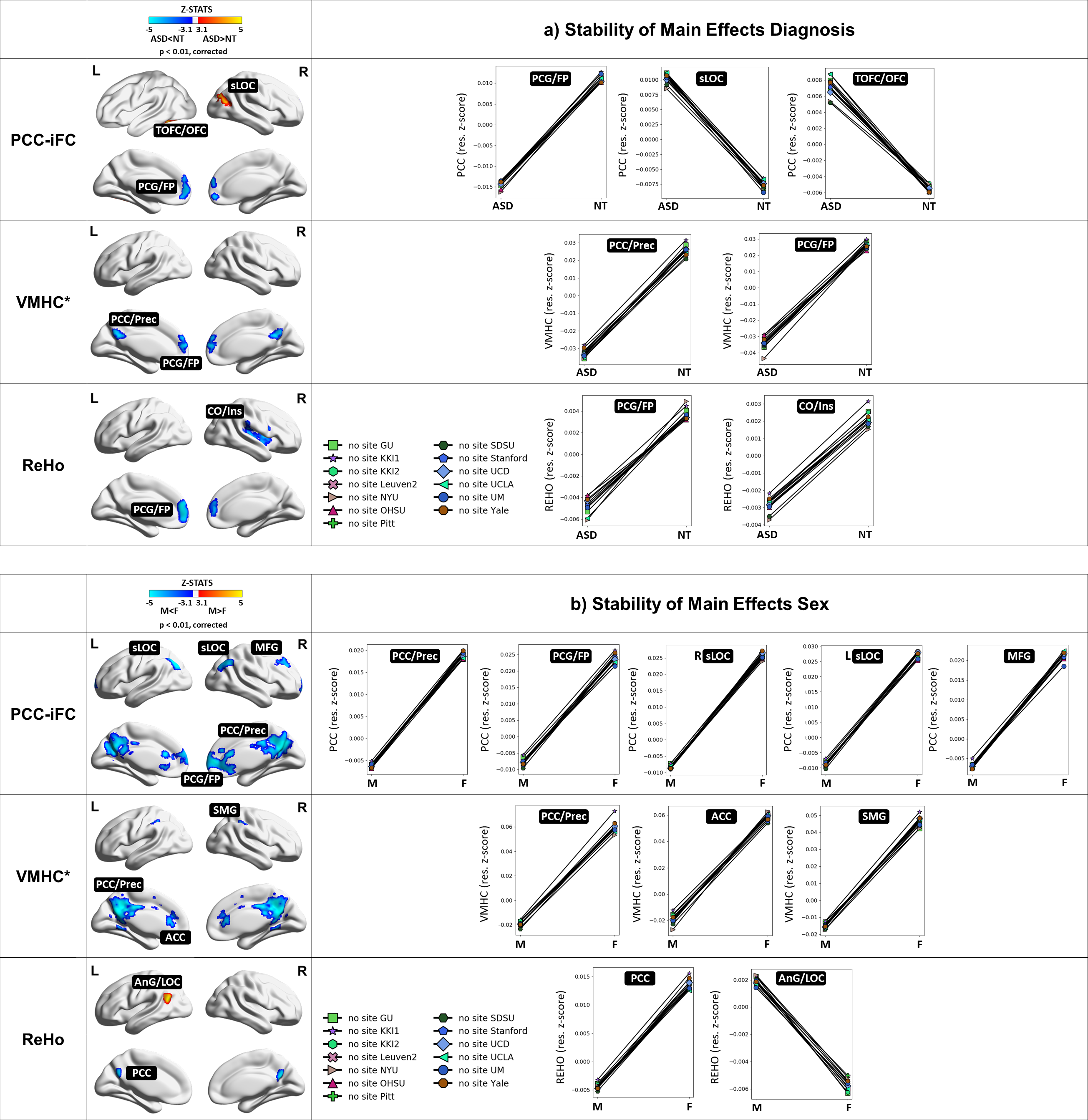

### Additional file 8

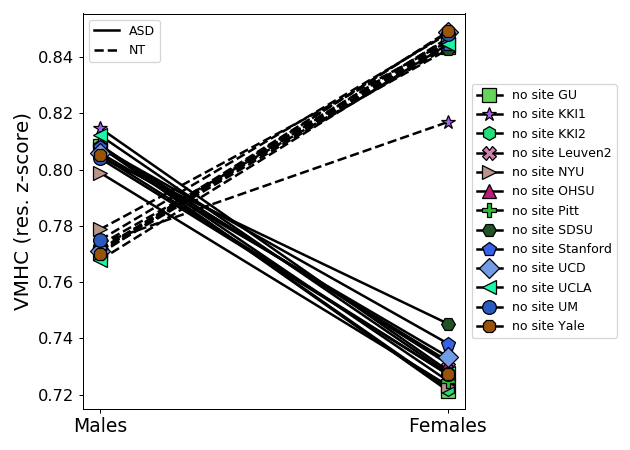

### Additional file 9

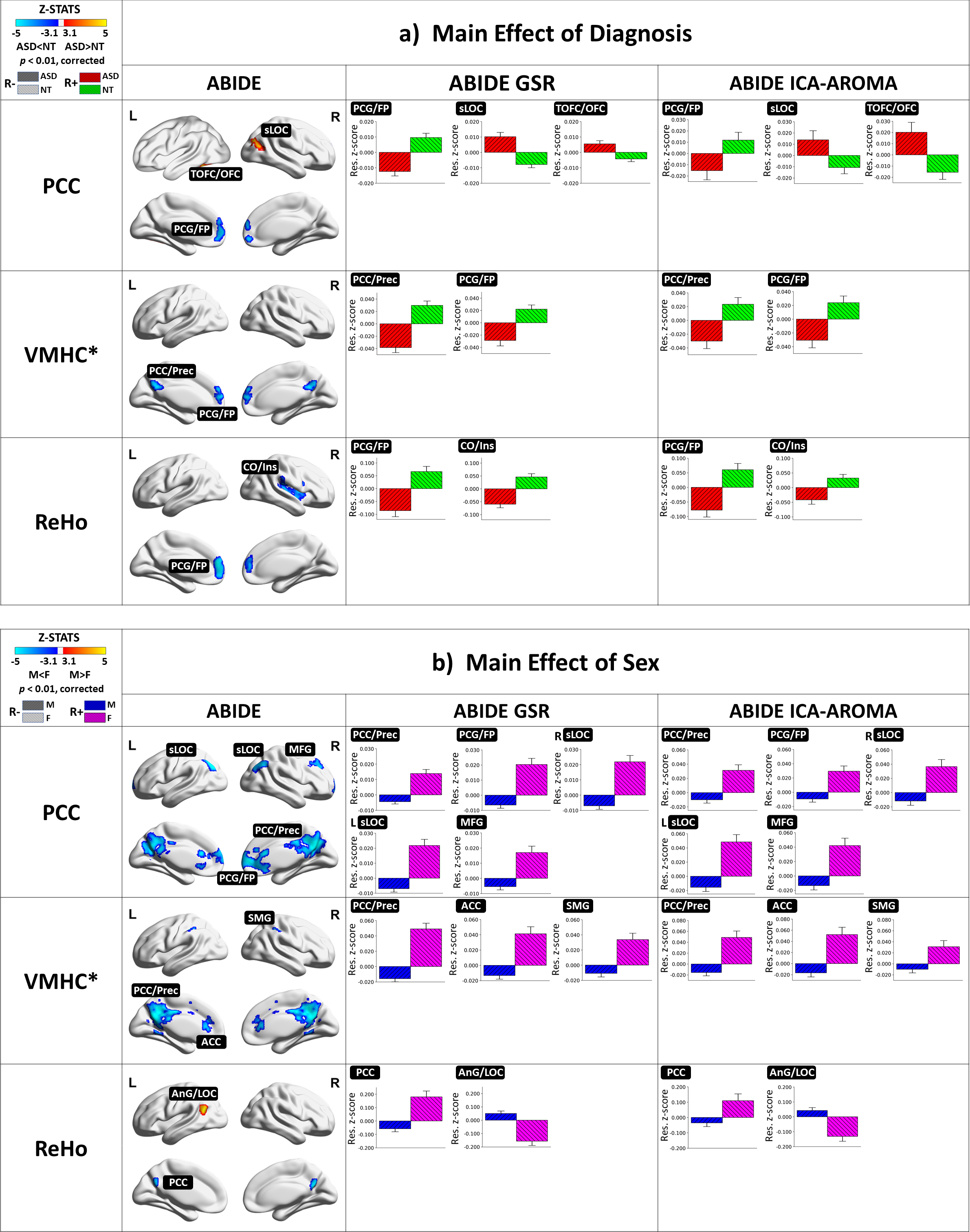

### Additional file 10

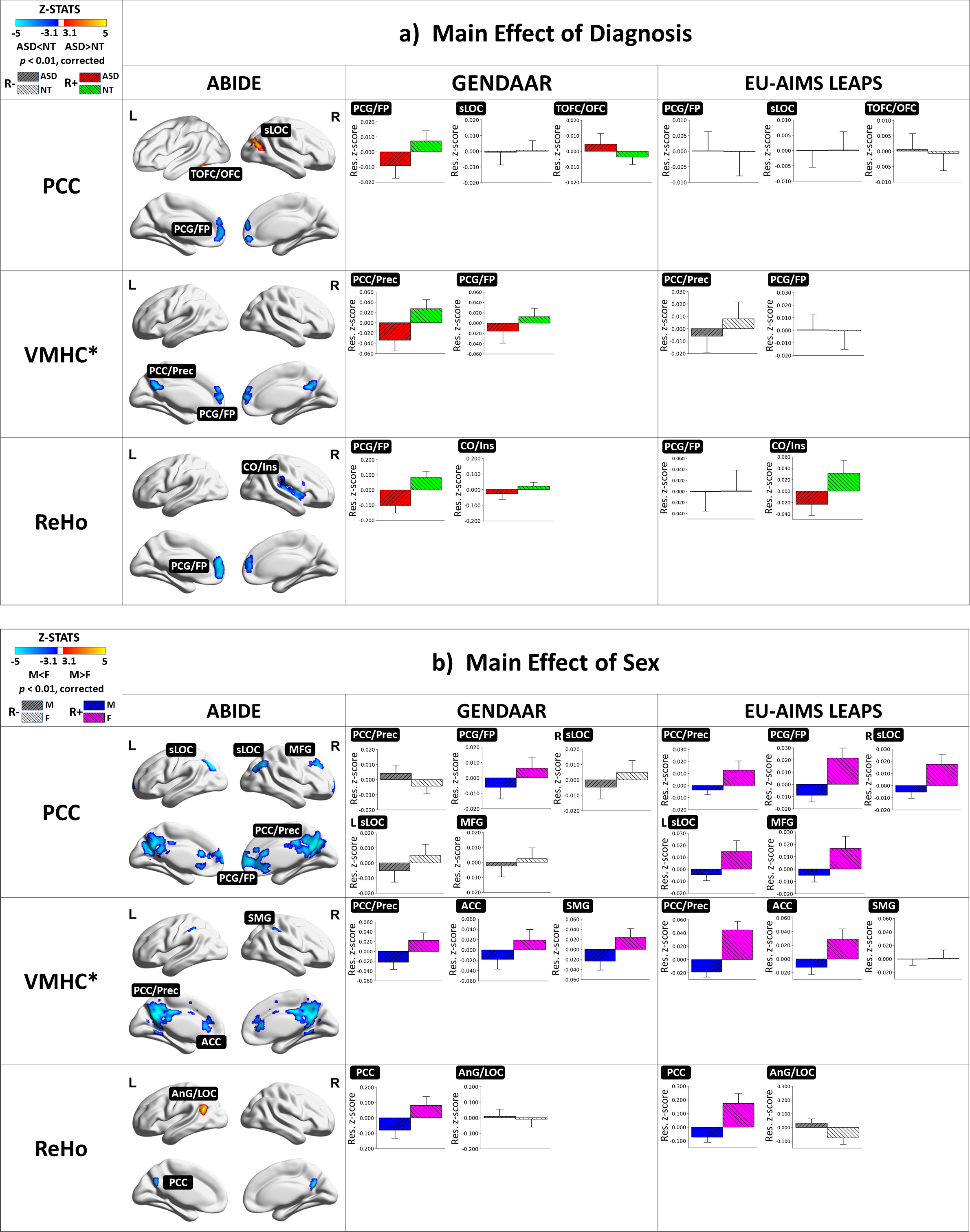
