## Additional file 5 for "Towards robust and replicable sex differences in the intrinsic brain function of autism"

| | | | | | | | DISCOVERY | | | ROBUSTNESS | | | | | | REPLICABILITY | | | | | |
|---|---|---|---|---|---|---|---|---|---|---|---|---|---|---|---|---|---|---|---|---|---|
| R-fMRI metric | Cluster | | Center of Gravity (MNI) | | | Statistic | CompCor (n=1019) | | | GSR (n=1019) | | | ICA AROMA (n=1019) | | | GENDAAR (n=196) | | | EU-AIMS (n=309) | | |
| | Anatomical Label | # of voxels | x | y | z | Z | $\eta_p^2$ | CI- | CI+ | $\eta_p^2$ | CI- | CI+ | $\eta_p^2$ | CI- | CI+ | $\eta_p^2$ | CI- | CI+ | $\eta_p^2$ | CI- | CI+ |
| **DIAGNOSIS** | | | | | | | | | | | | | | | | | | | | | |
| **PCC-iFC** | **PCG/FP** | 743 | -2 | 56 | 6 | 3.9 | 0.03 | 0.01 | 0.05 | 0.03 | 0.00 | 0.01 | 0.01 | 0.00 | 0.03 | 0.01 | 0.00 | 0.06 | 0.00 | 0.00 | 0.00 |
| | **sLOC** | 567 | 36 | 76 | -76 | 3.8 | 0.02 | 0.01 | 0.05 | 0.03 | 0.00 | 0.03 | 0.01 | 0.00 | 0.02 | 0.00 | 0.00 | 0.01 | 0.00 | 0.00 | 0.00 |
| | **TOFC/OFG** | 529 | -40 | -60 | -16 | 4.4 | 0.01 | 0.00 | 0.03 | 0.01 | 0.00 | 0.01 | 0.01 | 0.00 | 0.03 | 0.01 | 0.00 | 0.05 | 0.00 | 0.00 | 0.01 |
| **VMHC*** | **PCC/Prec** | 191 | -6 | -54 | 30 | 4.2 | 0.03 | 0.01 | 0.05 | 0.04 | 0.02 | 0.07 | 0.01 | 0.00 | 0.03 | 0.03 | 0.00 | 0.09 | 0.00 | 0.00 | 0.00 |
| | **PCG/FP** | 276 | -4 | 54 | 14 | 3.9 | 0.03 | 0.01 | 0.05 | 0.02 | 0.01 | 0.04 | 0.01 | 0.00 | 0.03 | 0.01 | 0.00 | 0.05 | 0.00 | 0.00 | 0.00 |
| **ReHo** | **PCG/FP** | 1072 | -2 | 54 | 10 | 5.2 | 0.01 | 0.00 | 0.03 | 0.02 | 0.01 | 0.05 | 0.02 | 0.01 | 0.04 | 0.04 | 0.00 | 0.11 | 0.01 | 0.00 | 0.05 |
| | **CO/Ins** | 1131 | 44 | -8 | 8 | 3.4 | 0.01 | 0.00 | 0.03 | 0.03 | 0.01 | 0.06 | 0.02 | 0.00 | 0.03 | 0.01 | 0.00 | 0.05 | 0.04 | 0.01 | 0.09 |
| **SEX** | | | | | | | | | | | | | | | | | | | | | |
| **PCC-iFC** | **MFG** | 676 | 28 | 32 | 42 | 3.4 | 0.03 | 0.01 | 0.06 | 0.03 | 0.01 | 0.05 | 0.02 | 0.01 | 0.04 | 0.00 | 0.00 | 0.03 | 0.01 | 0.00 | 0.04 |
| | **sLOC** | 901 | 46 | -66 | 42 | 5.2 | 0.04 | 0.02 | 0.07 | 0.03 | 0.02 | 0.06 | 0.04 | 0.00 | 0.03 | 0.00 | 0.00 | 0.04 | 0.03 | 0.00 | 0.08 |
| | **sLOC** | 966 | -40 | -74 | 40 | 6.7 | 0.05 | 0.03 | 0.08 | 0.04 | 0.02 | 0.07 | 0.03 | 0.01 | 0.05 | 0.00 | 0.00 | 0.04 | 0.03 | 0.00 | 0.07 |
| | **PCG/FP** | 2261 | 4 | 50 | 8 | 3.3 | 0.04 | 0.02 | 0.07 | 0.04 | 0.02 | 0.07 | 0.02 | 0.01 | 0.04 | 0.01 | 0.00 | 0.05 | 0.03 | 0.00 | 0.07 |
| | **PCC/Prec** | 4385 | 2 | -54 | 32 | 4.4 | 0.04 | 0.02 | 0.07 | 0.04 | 0.02 | 0.07 | 0.02 | 0.01 | 0.04 | 0.01 | 0.00 | 0.05 | 0.03 | 0.00 | 0.08 |
| **VMHC*** | **ACC** | 355 | -2 | 24 | 24 | 3.6 | 0.05 | 0.02 | 0.07 | 0.03 | 0.01 | 0.05 | 0.02 | 0.01 | 0.04 | 0.01 | 0.00 | 0.05 | 0.02 | 0.00 | 0.06 |
| | **SMG** | 404 | -48 | -36 | 44 | 3.6 | 0.04 | 0.02 | 0.06 | 0.02 | 0.01 | 0.04 | 0.01 | 0.00 | 0.03 | 0.02 | 0.00 | 0.07 | 0.00 | 0.00 | 0.01 |
| | **PCC/Prec** | 1785 | -8 | -48 | 30 | 4.5 | 0.07 | 0.05 | 0.11 | 0.07 | 0.04 | 0.10 | 0.03 | 0.01 | 0.05 | 0.02 | 0.00 | 0.08 | 0.06 | 0.02 | 0.12 |
| **ReHo** | **AnG/LOC** | 529 | -54 | -58 | 24 | 4.9 | 0.01 | 0.00 | 0.02 | 0.03 | 0.01 | 0.05 | 0.02 | 0.01 | 0.04 | 0.00 | 0.00 | 0.02 | 0.00 | 0.00 | 0.00 |
| | **PCC** | 444 | 2 | -58 | 26 | 4.3 | 0.04 | 0.02 | 0.06 | 0.03 | 0.01 | 0.05 | 0.01 | 0.00 | 0.03 | 0.02 | 0.00 | 0.08 | 0.01 | 0.00 | 0.05 |
| **SEX by DIAGNOSIS** | | | | | | | | | | | | | | | | | | | | | |
| **VMHC*** | ***sLOC*** | *138* | -30 | -78 | 28 | 3.7 | 0.02 | 0.01 | 0.04 | 0.01 | 0.00 | 0.03 | 0.01 | 0.00 | 0.02 | 0.00 | 0.00 | 0.03 | 0.01 | 0.00 | 0.03 |

*Abbreviations:* PCC-iFC: posterior cingulate cortex intrinsic functional connectivity, VMHC: voxel-mirrored homotopic connectivity, ReHo: regional homogeneity, TOFC/OFG: temporal occiptal fusiform cortex/occiptal fusiform gyrus, sLOC: superior lateral occipital cortex, PCG/FP: paracingulate cortex/frontal pole, PCC/Prec: posterior cingulate gyrus/precuneus, CO/Ins: central operculum/insula, MFG: middle frontal gyrus, ACC: anterior cingulate cortex, SMG: supramarginal gyrus, AnG/LOC: angular gyrus/lateral occipital cortex. *Due to processing failure of two subjects for VMHC, the sample size comprised 1017 subjects instead of 1019 for ABIDE and 307 instead of 309 for EU-AIMS. Color-code green: findings meeting criteria for robustness and replicability' yellow: findings not meeting criteria for robustness and replicability.
